# Non-random brain connectome wiring enables robust and efficient neural network function under high sparsity

**DOI:** 10.64898/2026.03.30.715411

**Authors:** James McAllister, Conor Houghton, John Wade, Cian O’Donnell

## Abstract

The connectivity of brain networks is extremely sparse due to metabolic, physical and spatial constraints. Although wiring sparsity can confer computational advantages for biological and artificial neural networks, sparse networks require fine parameter tuning and exhibit strong sensitivity to perturbations. How brains achieve their efficiency and robustness is unclear. Here we addressed this by analysing the dynamical properties of Echo State Networks with wiring based on the *Drosophila melanogaster* fruit fly connectome, compared with sparsity-matched random-wiring networks. We evaluated these networks on a set of eight cognitive tasks, and found that connectome-based neural networks (CoNNs) typically showed narrowly distributed task engagement across their neurons. The importance of a neuron for task performance correlated with its node degree, local clustering, and self-recurrency, and these correlations were stronger in CoNNs than in random networks. CoNNs were more robust to neuronal loss, retaining their task performance and beneficial dynamical properties such as criticality and spectral radius better than random networks. Similarly, CoNNs were more robust to hyperparameter variations in both input and recurrent weight scaling. Using theoretical arguments and numerical simulations, we show that excess CoNN node self-recurrency is sufficient to explain this enhanced robustness. Overall, these results identify non-random features of connectome wiring that allow brains to reconcile extreme sparsity with reliable computation.

**Significance:** Brain networks support robust computation even though they operate under extreme wiring sparsity due to metabolic and spatial constraints. While sparse networks typically require fine-tuning and are sensitive to perturbations, we show that biological connectomes support specialised, efficient task engagement and remain robust to neuron loss and parameter variation. We identify excess neuronal self-recurrency as a key structural feature underlying this stability. These results reveal how non-random connectivity stabilises computation in extremely sparse networks, providing principles for understanding brain function and designing robust, efficient artificial neural systems.

## Introduction

Brain network connectivity is extremely sparse, with only a tiny fraction of all possible neuron pairs actually forming synaptic connections. This sparsity is ubiquitous across organisms (1, 2), and there are various factors explaining why circuit development results in such sparsity, including energy costs, and physical or spatial constraints (3, 4). In addition to optimising energy efficiency, extreme sparsity is also an advantage for computational efficiency in learning, information processing, and pattern separation (5–7). Network sparsity is not only an important feature in biological contexts, but sparse artificial neural networks in machine learning and neuromorphic computing yield functional benefits: reduced power consumption, computational cost, storage requirements, and improved training speeds, generalisability, and scalability (8, 9).

However, extreme sparsity in neural networks can come at the cost of a reduction in robustness: sparse networks need to be carefully fine-tuned and they are highly sensitive to perturbations (10–12). Given that biological network development is noisy (13, 14) and yet extremely sparse, the question is: how do brains achieve robust and stable computation under such extreme sparse wiring constraints? We used computational modelling of Echo State Networks (ESNs) based on biological brain connectivity to explore this question.

ESNs are a type of reservoir computer (15, 16). Unlike traditional recurrent neural networks (RNNs) where all the synaptic weights are changed during training, in an ESN the recurrent core is fixed. This core is typically a random network which produces rich transformations of the input. Network activity is mapped to a readout and only these readout weights are trained by optimisation (Figure 1A). This framework enables implementing specific network topologies or features of interest within the RNN, to analyse resulting dynamics and performance. Due to their theoretical tractability and practical success in tasks such as short-term memory, time-series prediction, classification, and dynamical system modelling (17–20), ESNs are a powerful tool for studying the interplay between network structure and dynamics.

**Figure 1.**
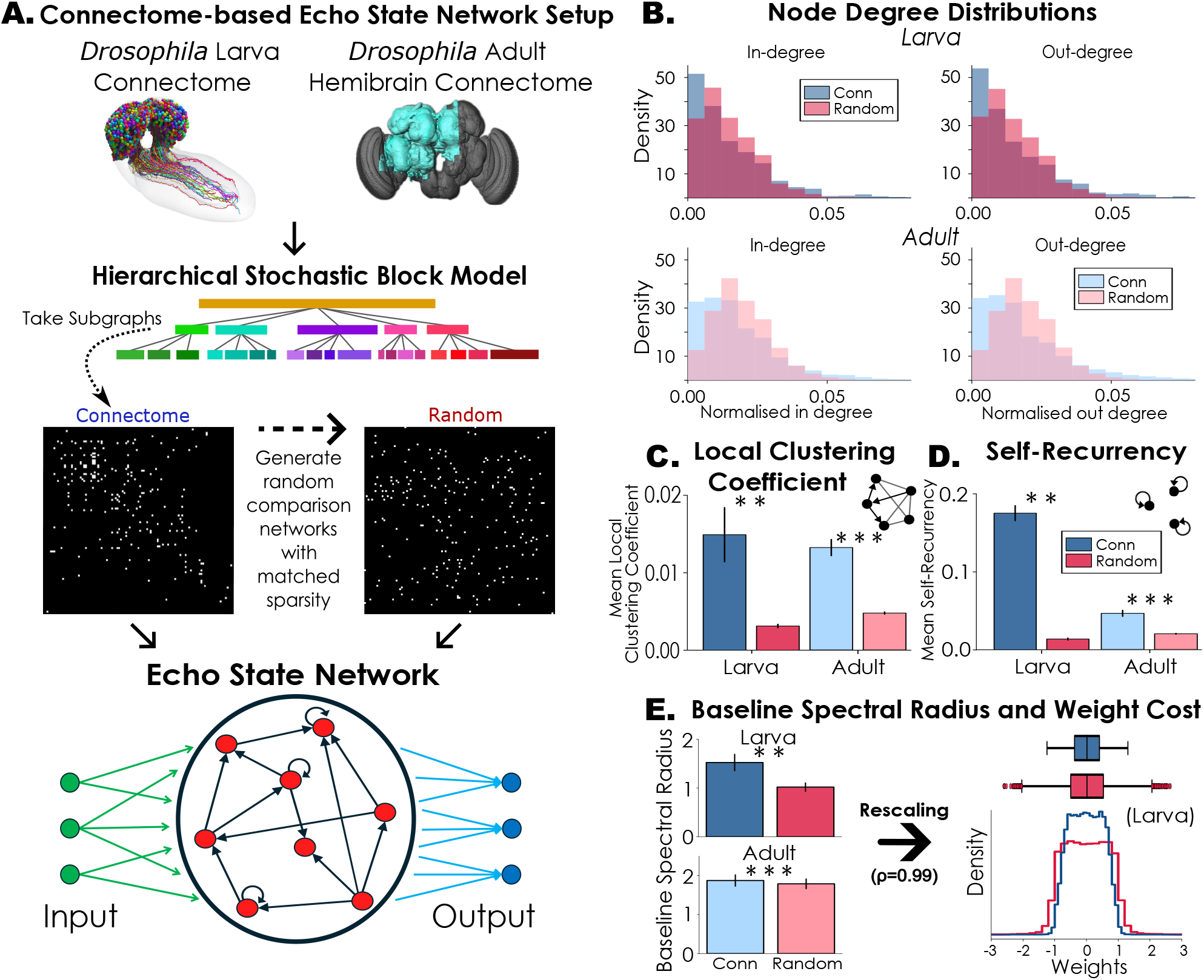
**A Illustration of the connectome-based Echo State Network (CoNN) setup.** A hierarchical stochastic block model detection algorithm was applied to the larval (21) and adult (22) *Drosophila melanogaster* connectomes, and subgraphs were extracted. Equivalent random networks were generated for comparison. Edge weights were drawn uniformly between *−*1 and 1, and then the weight matrices were rescaled to achieve a desired spectral radius. These connectome-based and random networks were then used in Echo State Networks. **B–D. Network Feature Comparisons between connectome and random networks. B**. Node in (left) and out (right) degree distributions for larva (top) and adult (bottom) networks. CoNN statistics shown in blue, random networks in red. **C–D**. Mean local clustering coefficient (C) and mean self-recurrency (D). **E. Baseline Spectral Radius and Wiring Cost**. When weights are drawn from the same distribution, the baseline spectral radius is higher for CoNNs than random ones (left). After rescaling to equivalent spectral radii, CoNNs have a narrower distribution of weights, and therefore a lower “wiring cost” (larva example shown on the right). Bars indicate standard error of the mean. (* * p < 0.01, * * * p < 0.001)

The ESN framework is a possible model for brain function (23), and so biological inspiration has lead to a focus on how the structure of the core RNN can be quantified, adjusted and compared to brain connectivity (24– 34). This has culminated in studies implementing ESNs with connectivity based on the synapse-resolution connectome of the the fruit fly (35, 36), where the central idea is that detailed biological connectivity embodies structural principles that may confer computational advantages over purely random networks. In chaotic timeseries prediction tasks connectome-based ESNs have smaller performance variance and often better average performance than classic random ESNs (37), with a robustness to hyperparameter variations (35). Connectome ESNs also exhibit greater capacities for multifunctionality across broader hyperparameter ranges than random networks (38). Multifunctionality is the ability to perform multiple mutually exclusive tasks without altering the network connections. While these results indicate a degree of versatility and robustness in biologically-based neural networks, it remains unclear how — and to what extent — connectome-based networks achieve functional robustness and efficiency in the face of extreme connectivity sparsity.

Here we address these gaps by implementing connectome-based neural networks (CoNNs) from larval (21) and adult *Drosophila melanogaster* (22) connectomes as ESNs (Figure 1A). These ESNs were tested in in tasks involving memory, decision-making, time-series prediction, and chaotic time-series predic-tion, and in analyses of structure and dynamics. CoNNs exhibited pronounced non-random network features (e.g. heterogeneous node degree distributions, local clustering, and self-recurrency), and, for matched initial weight distributions, yielded higher baseline spectral radii than random null models. After rescaling to an equivalent spectral radius, this resulted in a lower total “weight cost” in CoNNs.

In working memory, decision-making and time-series prediction tasks, CoNNs achieved comparable performance to classic random networks, and often outperform them when normalised for wiring cost. Using a *weighted task variance* framework, we show that CoNNs displayed more specialised and sparse patterns of task-relevant neural engagement, with performance that is more robust to both random and targeted node pruning. We identified relationships between neural contribution and local network features such as self-recurrency, node degree, and clustering coefficient, as well as biological cell-type annotations.

We developed theoretical scaling approximations for the network spectral radii, demonstrating how sparsity and self-recurrency influence spectral radius and its robustness to network pruning. CoNNs showed greater robustness of spectral radius and maximum Lyapunov exponent to both random and targeted node pruning. Similarly, CoNNs displayed a greater robustness of dynamical properties in hyperparameter variation, albeit with indications of lower-dimensional activity.

## Results

### Network Features

The connectivity of biological neural networks is extremely sparse. For example, the mean connectivity sparsities (the fraction of absent connections) of the larval and adult *Drosophila* networks used in this study were 0.9859 and 0.9802, respectively. In contrast, ESN networks are commonly initialised with a connectivity sparsity around 0.8 – 0.9 (39, 40, 18, 19, 30), corresponding to 5–10-fold more connectivity than the connectomes.

To obtain reduced networks that preserve biologically meaningful structure (41) we partitioned the full networks into communities using a hierarchical stochastic block model (42) (Figure 1A middle). We selected small subnetworks (sizes 100 to 400 neurons) and built ESNs using their connectivity graphs. As control comparisons, we generated random networks (Erdős-Rényi graphs) with matched sparsities to those from the connectome (Figure 1A). These random networks lack specific network structures — such as clusters, block structure, enhanced recurrency, or heterogeneous node degree distributions — which may be present in trained, engineered, or biologically-developed networks. For an alternative comparison, we also generated equivalent Configuration Model networks, which matched the degree distributions of the connectome networks, but otherwise randomised all wiring, eliminating other higher-order structural features. These Configuration Model networks generally displayed similar results to the random Erdő s-Rényi networks used throughout this study (Supplementary Material).

To characterise basic network features of connectomes, we first calculated the node degree distributions, mean local clustering coefficient, and mean self-recurrency in both the connectome-based and random networks. Figure 1B–D shows how these three network features differ significantly between connectome and random networks. Figure 1B shows that the distributions of connectome degrees are more heterogeneous and heavy-tailed (43, 44) than their random counterparts. The connectome-based networks (CoNNs) also show significantly higher degrees of local clustering (*p* = 0.004 and *p* < 0.001 for larva and adult, respectively) and self-recurrency (*p* = 0.004 and *p* < 0.001 for larva and adult, respectively) than random models (Figure 1C and D). Larval CoNNs are also more self-recurrent than the adult CoNNs (*p* < 0.001, Mann-Whitney U Test).

### Spectral Radius and Wiring Cost

The spectral radius *ρ* of a network is the largest absolute eigenvalue of its recurrent weight matrix, and it plays a significant role in the network dynamics. In random networks, the spectral radius controls the transition between contractive, stable dynamics and expansive or chaotic regimes, and is therefore a key parameter in ESN computational performance (15, 18, 45). For random matrices with i.i.d. weights, random matrix theory predicts 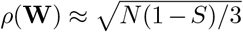 ((46) and Supplementary Material 1), showing that baseline spectral radius depends on network size and density.

In sparse heterogeneous networks, however, the spectral radius scales approximately with the square root of the maximum degree (47, 48). Consistent with this, both larval and adult CoNNs exhibit significantly larger baseline spectral radii than their random null models (Figure 1E left and S1; *p* = 0.004 and *p* < 0.001 for larva and adult, respectively).

To ensure comparable dynamical regimes, we rescaled all weight matrices to a common spectral radius. Because CoNNs start with larger baseline spectral radii, rescaling results in smaller effective weights relative to random networks (Figure 1E right and S1). Consequently, the total absolute weight (a proxy for wiring cost (3)) is systematically lower in CoNNs.

### Task Performances and Costs

We compared the performances of the ESNs on eight computational and cognitively relevant tasks, arranged in four pairs: 1a. Memory Capacity, 1b. Sequence Recall, 2a. Two Input Decision-Making, 2b. Delayed Decision-Making, 3a. Oscillator Time-Series Prediction, 3b. Lotka-Volterra Time-Series Prediction, 4a. Lorenz System Chaotic Time-Series Prediction, and 4b. Rössler System Chaotic Time-Series Prediction. Over-all, CoNNs performed comparably to random networks (Figure S16A), except for the Memory Capacity and Sequence Recall tasks.

Biological networks may optimise a trade-off between computational capacity and wiring cost (33). To account for the “wiring cost” of the networks, we normalised the task performances of each ESN by the sum of the absolute values of the network weights. Following this normalisation, CoNNs outperformed the conventional ESNs across most tasks (Figure S16B).

To characterise a biological proxy of “energy cost” of the networks during tasks, we quantified an activity-based measure by computing, for each task, the total squared neural activity (i.e. the total squared deviation from in-activity). Across all tasks, CoNNs exhibited lower energy costs than random networks (Figure S17), indicating that connectome task performances were achieved with reduced overall state magnitudes.

### Distributions of Neural Contribution

To capture the level of engagement and contribution of a neuron during a given task, we introduced a measure called *weighted task variance* (WTV), capturing the idea that neurons that contribute more to a given task should have both time-varying activity and large readout weights (Methods, (49)). Then to measure the spread of neural engagement across tasks and across networks, we calculated the mean normalised participation ratio of the WTV (i) for each neuron across tasks, and (ii) for each task across neurons. These two approaches addressed (i) how task-specialised or generalised the neurons are, and (ii) how concentrated or distributed the neural engagement is (Figure 2A).

**Figure 2.**
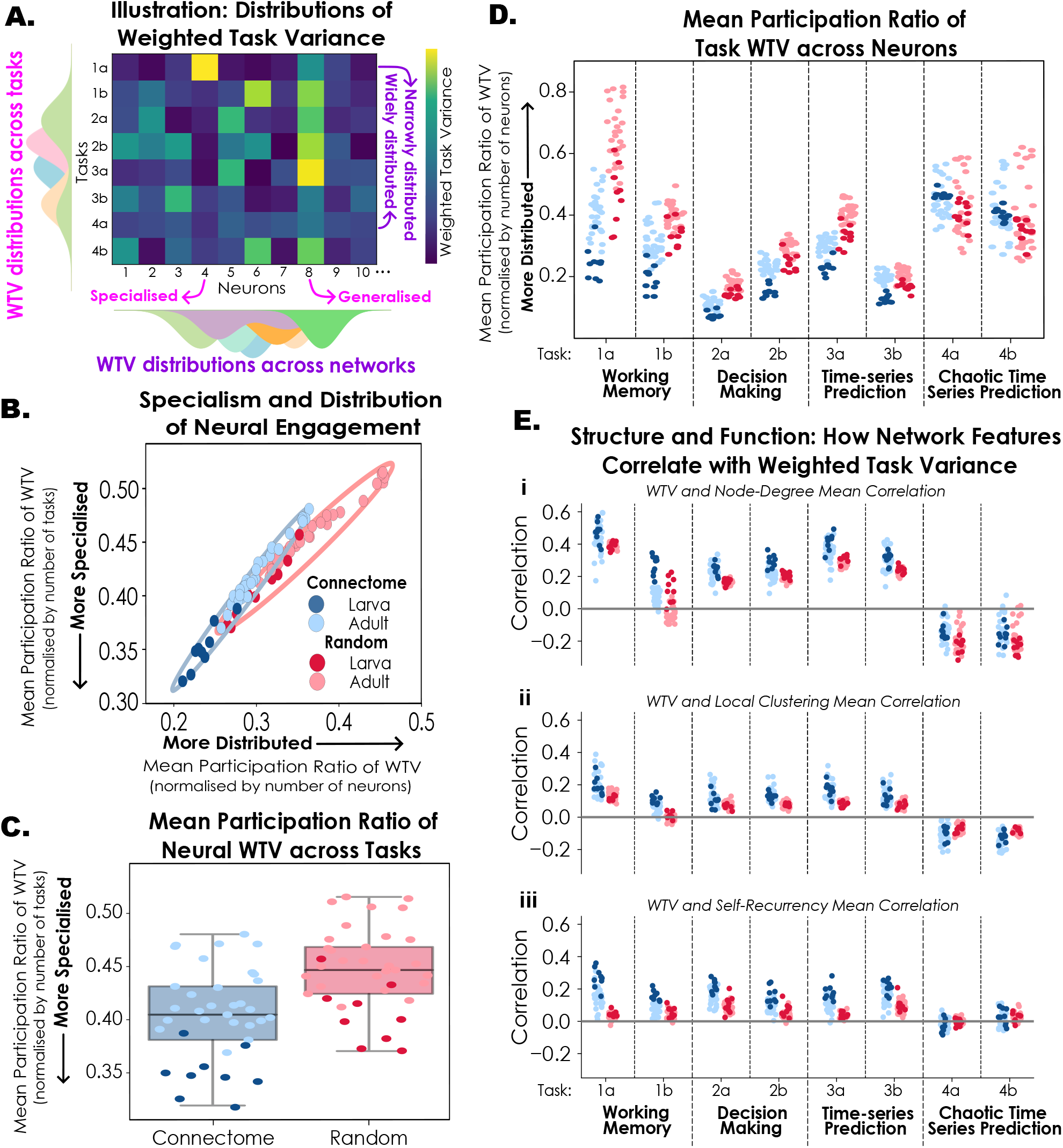
A–D. Task Specialisation and Distribution of Neural Engagement. **A**. Illustration of how neural task variance distribution and specialisation are calculated from the weighted task variance across all neurons and all tasks. The WTV of Neuron 4 is specialised for Task 1a, but for Neuron 8 it is generalised across several tasks. Task 1a WTV is narrowly distributed across few neurons, but for Task 4a it is widely distributed across neurons. **B**. Mean participation ratio of weighted task variance in 2D space: Specialism and Distribution of Neural Contribution. (Ellipses show the minimum-volume enclosing ellipses of the connectome and random points). **C**. Distribution of Neural Contribution across tasks: Mean participation ratio of weighted task variance for each neuron across 8 tasks — measures how task-specialised or generalised each neuron is. **D**. Distribution of Neural Contribution during each task: Mean participation ratio of weighted task variance across neurons for each task — measures how narrowly or widely distributed the neural contribution is. **E. Structure and Function: How network features correlate with weighted task variance**. Mean correlations between WTV and network features of (i) node degree, (ii) local clustering coefficient, (iii) self-recurrency.

CoNNs had a lower mean task WTV participation ratio averaged across neurons (Figure 2C) than random networks (*p* = 0.004 and *p* < 0.001 for larva and adult, respectively), indicating a higher degree of task specificity, whereas the mean task participation ratio of equivalent random networks was higher and therefore more generalised. Second, the mean network WTV participation ratio was lower for CoNNs than for random networks on Tasks 1–3. (*p* = 0.004 for larva and *p* < 0.001 for adult). This suggests that a smaller subset of neurons in the network was engaged and contributing to these tasks.

The same was not observed in the chaotic time-series prediction tasks (Tasks 4a and 4b), where the CoNNs had higher mean network WTV participation ratio than random networks (*p* = 0.004 and *p* = 0.008 for larva in tasks 4a and 4b respectively, and *p* = 0.032 for adult in Task 4a but not significantly different for adult in Task 4b, *p* = 0.099). The Configuration Model networks generally showed intermediate WTV participation ratios between the connectome and random Erdő s-Rényi networks, across both tasks and networks (Figure S25B and C).

In Tasks 1–3 larval CoNNs showed a lower mean network WTV participation ratio than adult networks (*p* < 0.001, Mann-Whitney U tests), indicating a greater degree of neural specialism in larva vs adult.

### Network Structure and Function

To test if local network structure can account for differences in WTV between connectome and random networks, we computed the correlation between WTV and select network features: node degree, local clustering coefficient, self-recurrency, and biological cell type.

Across Tasks 1–3, the network features of node degree, local clustering coefficient, and self-recurrency were positively correlated with weighted task variance (Figure 2E); this suggests that the more important a neuron is for these tasks, the more likely it is to have a higher node degree, higher local clustering coefficient, or be self-recurrent. In contrast, the chaotic time-series prediction tasks (Tasks 4a and 4b) showed either very weak or negative correlation between these structural properties and weighted task variance. These qualitative trends — while stronger for the CoNNs — were present for both connectome and random networks (blue vs red Figure 2E).

To test if there is a relationship between neural engagement and biological cell type, we correlated WTV with previously published cell type annotations (21, 22), then averaged all neurons of a given cell type, and averaged over all 9 larval and 28 adult networks. For Tasks 1–3, we found heterogeneity in cell type engagement, whereas for Tasks 4a and 4b, all cell types participated approximately equally, both for larval (Figure S26) and adult (FigureS27) CoNNs. This is consistent with the results for the participation ratio of WTV (Figure 2D). The adult networks tended to show more structured cell-type task specialisation than larval networks (Supplementary Note 9).

### Pruning Neurons

Pruning nodes from neural networks is commonly used as a way to 1) model neuronal loss or damage for both biological and artificial neural networks (50, 51); 2) reduce artificial network sizes (52, 9, 11); 3) test causal neural contributions to function (53). Here, we tested the effect of node pruning on ESNs task performance, spectral, and dynamical properties.

#### Pruning and Task Performance

To test how pruning affects task performance, we removed neurons from each network in order from least to most important, as captured by their weighted task variance, retraining the output layer each time. Task performance generally decreased with increasing node removal, however the CoNNs were typically more robust, showing slower performance degradation compared to random networks (examples in Figure 3A). To summarise this effect, we asked what fraction of the network needs to be pruned to cause a 10% relative performance drop. CoNNs were either as robust or more robust than random networks for all 8 tasks (Figure S18).

**Figure 3.**
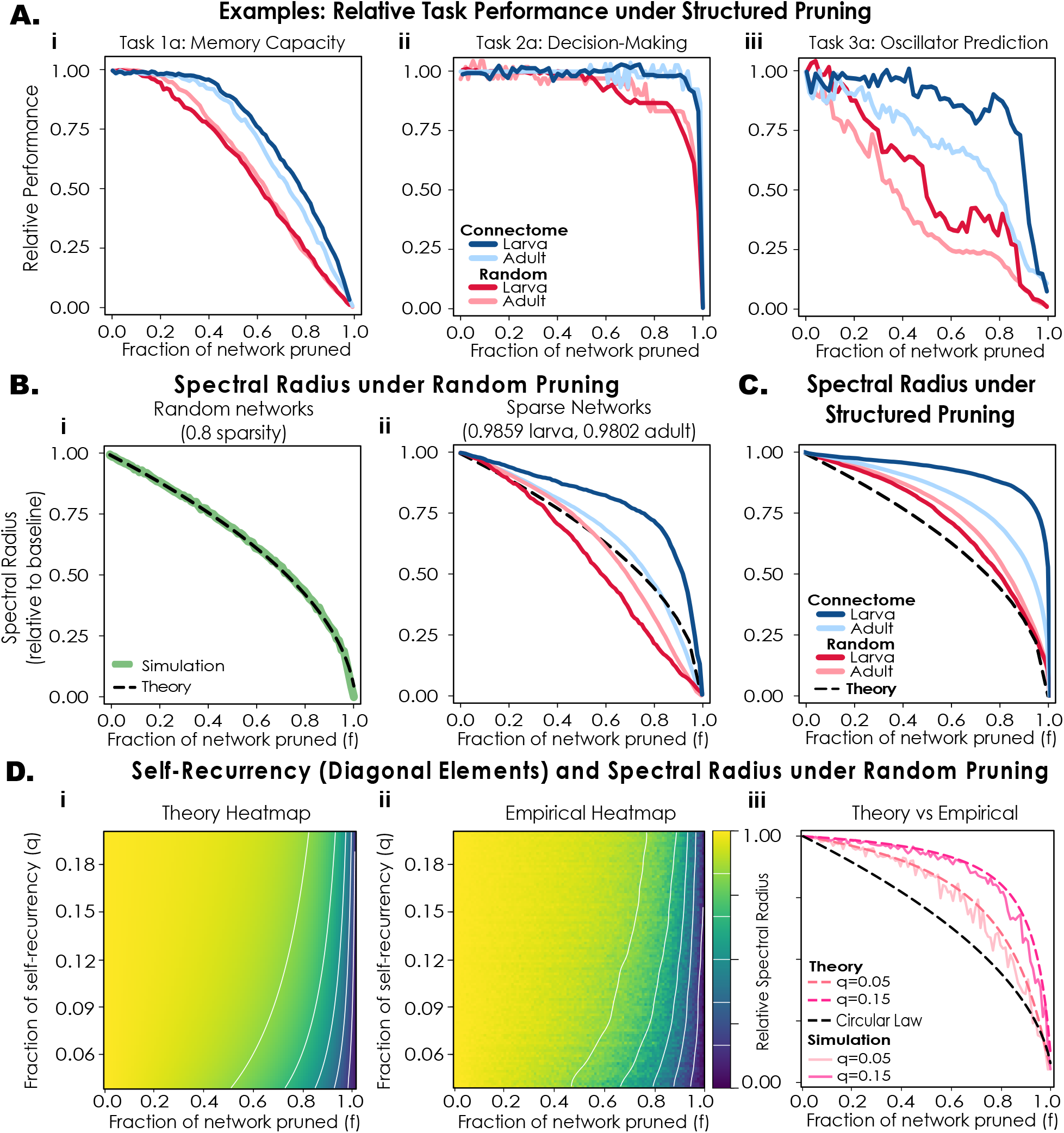
**A Example relative task performance under structured pruning.** (i) Task 1a: Memory Capacity, (ii) Task 3a: Decision-Making and (iii) Task 5a: Oscillator Prediction relative task performances as neurons are pruned in order of increasing weighted task variance. **B. Spectral radius and random pruning**. i. Circular Law prediction from Random Matrix Theory and simulations of relative spectral radius over random pruning. ii. Random pruning of the sparse ESNs in this study, compared with the Circular Law prediction. **C. Spectral radius and structured pruning**. Relative spectral radius as neurons are removed in order of increasing weighted task variance (averaged across tasks), and compared with the Circular Law prediction. **D. How self-recurrency (non-zero diagonal elements) impacts on relative spectral radius under random pruning**. i. Theory-based heatmap showing how the fraction *q* of non-zero diagonal elements stabilises spectral radius decrease over random pruning (network size *N* = 100, sparsity *S* = 0.985). White contour lines trace out equal values of relative spectral radius. ii. Empirical heatmap as in (i) for 100 trials. iii. Example relative spectral radius under random pruning for different self-recurrency fractions *q*, showing how rearranging elements from sparse random matrices onto the diagonal impacts relative spectral radius robustness.

#### Pruning and Spectral Radius

To examine how neuronal pruning affects network spectral radius, removing *m* neurons from a random network of size *N* and sparsity *S* produces an (*N* −*m*) × (*N* −*m*) sub-matrix whose entries remain independent with sparsity *S* and variance *σ*^2^. By the Circular Law

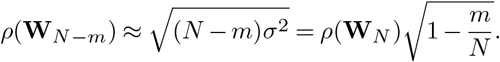

Hence, the spectral radius is expected to scale as 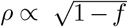, where *f* = *m/N* is the fraction pruned. This is a useful baseline: any deviation from this scaling under random pruning reflects important non-random network organisation. Although derived in the dense, infinitesize limit, we tested this scaling by randomly pruning 100 networks size *N* = 100, sparsity *S* = 0.8. Figure 3Bi shows good agreement with the 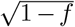 prediction even in this finite, moderately sparse regime.

However, when we randomly pruned the connectome and random networks in this study, something different emerged. The spectral radius of random networks fell more rapidly than Circular Law prediction, particularly in the sparser larva-comparison networks (Figure 3Bii). In contrast, the spectral radius of CoNNs declined slower than Circular Law prediction, particularly in the larva case, reflecting a non-random topology that induces robustness to pruning.

We also pruned neurons from the networks in order of weighted task variance, from least to most important, and averaged across networks and tasks. Under this structured pruning, the CoNNs again decreased in relative spectral radius more slowly than random networks (Figure 3C); this spectral decrease is noticeably slower than in the random pruning case (Figure 3Bii). The relative spectral radius of Configuration Model networks under both random and structured pruning more closely followed the random Erdős-Rényi networks than the CoNNs (Figure S25D), suggesting that the connectome’s node degree distribution by itself does not induce robustness to node removal.

A measure of network dynamics related to spectral radius is the maximum Lyapunov exponent (MLE). The MLE provides a measure of dynamical stability and sensitivity to perturbations. We computed the MLE (54) under both random and structured pruning, to see how node removal affects the dynamics of the networks. The MLE closely mirrored the spectral radius behaviour: CoNNs retained greater robustness to pruning than their random counterparts (Supplementary Figure S19).

#### Self-Recurrency and Spectral Radius Robustness

To better understand how self-recurrency (which is prevalent in the connectome) affects spectral radius robustness under pruning, we derived a theoretical approximation that accounts for diagonal and off-diagonal contributions to the spectral radius in sparse random networks. Using intuition from the Gershgorin Circle theorem ((55) and Supplementary Material 5), we approximated the spectral radius as the dominant contribution between the diagonal and bulk components of the matrix spectrum. Using extreme value statistics and the Circular Law from random matrix theory, we approximated the predicted relative spectral radius of a matrix with uniformly distributed weights by

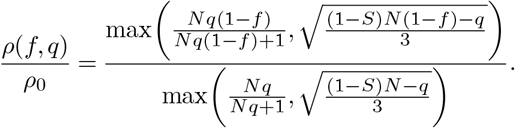

where *N* denotes the network size, *S* sparsity, *q* the fraction of self-recurrent neurons, and *f* the fraction of the network pruned. This approximation captures the competition between diagonal structure (self-recurrency) and the random bulk spectrum (Supplementary Information 5). For relatively small and sparse networks, this predicts that increasing the proportion of diagonal elements will lead to the diagonal contribution dominating the spectral radius.

We first visualised the theoretical scaling across different values of *q* (from 0.04 to 0.2) for networks of size *N* = 100 and sparsity *S* = 0.985 (Figure 3Di). Increasing the fraction of self-recurrent neurons systematically slows the decay of the relative spectral radius under pruning.

To test this prediction empirically, we generated sparse random networks while explicitly controlling both the overall sparsity and the diagonal fraction *q*. For each network, we first generated a directed random matrix (size *N* = 100, sparsity *S* = 0.985), and then reassigned a subset of existing off-diagonal weights onto the diagonal such that a fraction *q* (from 0.04 to 0.2) of neurons possessed non-zero self-connections while preserving the overall sparsity. The networks were then subjected to random node pruning and the spectral radius recomputed after each pruning step. The resulting empirical relative spectral radii were averaged across 100 simulations (Figure 3Dii). Simulations closely matched the theoretical predictions: increasing self-recurrency systematically enhanced spectral robustness under pruning. Theoretical and empirical pruning curves for representative values *q* = 0.05 and *q* = 0.15 are shown in Figure 3Diii. In both cases the theoretical approximation closely tracked the simulated spectral radius decay, confirming that the balance between diagonal extreme-value statistics and the random bulk spectrum provides a simple explanation for the stabilising effect of self-recurrency. These results suggest that high degrees of self-recurrency in conjunction with random sparse connectivity may serve as a mechanism for spectral robustness, potentially providing functional stability against random node loss.

## Dynamics and Dimensionality

To understand how dynamical capacities differ between connectome-based and random networks, we computed the maximum Lyapunov exponent and Lyapunov dimension of both across varying recurrent weight scaling (spectral radius in [0.01, 20]) and input layer weight scaling (in [0, 20]). These dynamical measures are important determinants of network capabilities for stability, responsiveness, and information processing (54, 28, 56). We found that the input weight scaling did not play as significant a role as spectral radius in defining dynamics.

Unlike in random dense networks (28, 57), we did not observe a ubiquitous transition to chaos (MLE *>* 0), with most networks remaining stable or at criticality for all the values of spectral radius. This is a consequence of the extreme sparsity of the connectivity matrices, which limits recurrent amplification. Supplementary Figure S23 demonstrates how lowering sparsity leads to more chaotic transitions.

Larva CoNNs remained near criticality with MLE values close to 0 over a broader range of spectral radii, compared with random networks which exhibited a narrower window of criticality (Figure 4Ai left). This suggests that CoNNs are more robust to the effects of recurrent weight scaling and spectral radius variation, preserving critical dynamics which are important for computation (58–61). Adult networks (Figure 4Aii left) did not show distinct differences between connectome and random networks.

**Figure 4.**
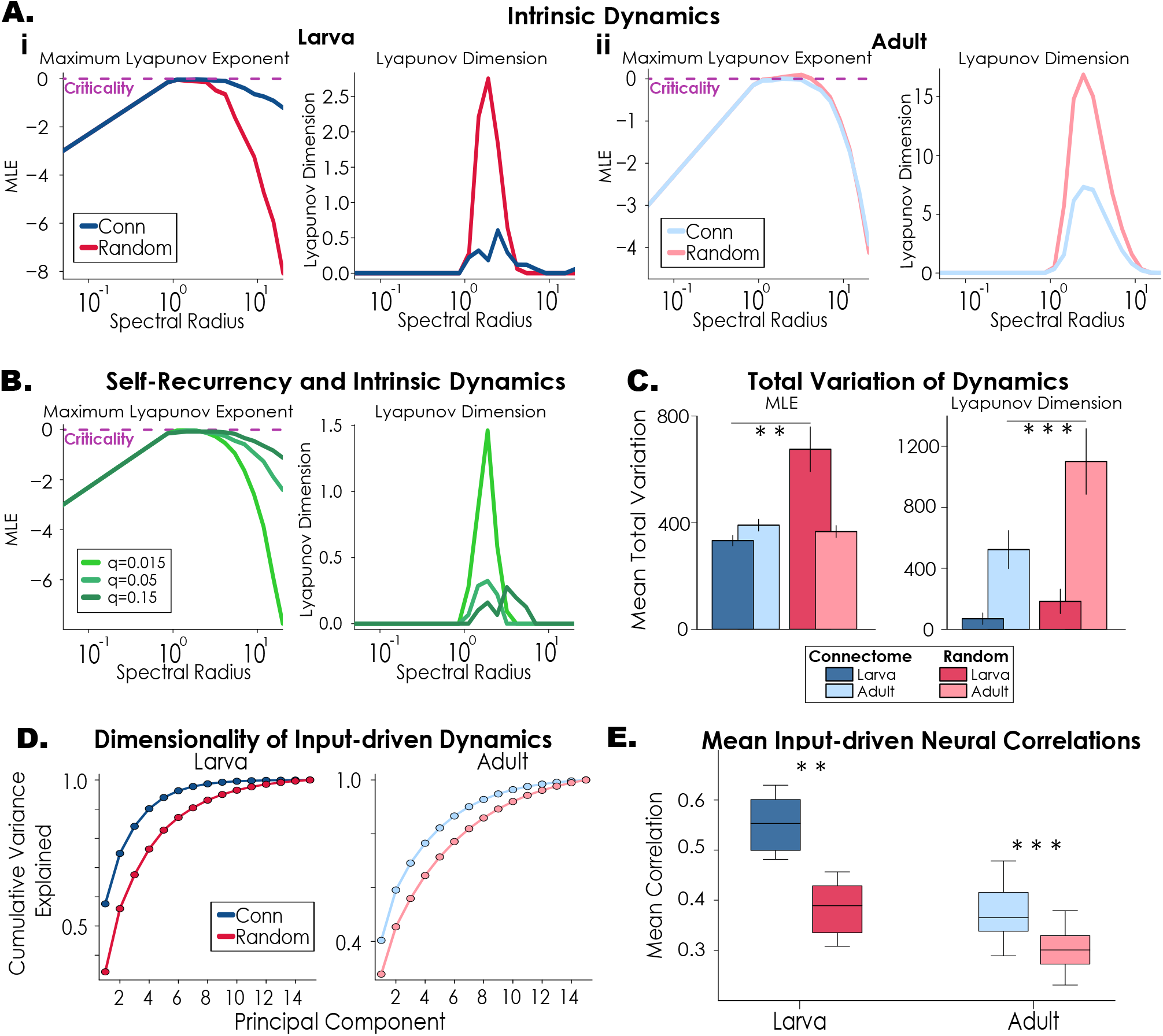
Dynamics and Dimensionality. **A**. The mean MLE (left) and Lyapunov dimension (right) across a range of spectral radius values for (i) larva and (ii) adult comparisons. **B**. Mean MLE (left) and Lyapunov dimension (right) across a range of spectral radius values for networks with varying degrees of self-recurrency. **C**. Mean total variation of MLE and Lyapunov dimension for connectome and random networks (bars show standard error of the mean). The lower the total variation, the smoother the dynamical landscapes. **D**. Dimensionality of network dynamics (PCA of network activity) driven by a noisy input, measured by cumulative variance explained. **E**. The mean absolute pairwise correlations of neural dynamics while being driven by a noisy input. (* * p < 0.01, * * * p < 0.001)

To investigate whether the higher self-recurrency in the larva CoNNs plays a role in maintaining robust critical dynamics under parameter variation, we generated random networks (size *N* = 100, sparsity *S* = 0.985) with differing fractions of self-recurrent neurons: *q* = 0.015 (baseline), 0.05 and 0.15. We computed MLE and Lyapunov dimension of these networks over a range of spectral radii in [0.01, 20]. Increasing self-recurrency lead to dynamics remaining closer to criticality for a wider range of spectral radii (Figure 4B left), consistent with with the larva CoNNs.

To quantify how smooth or fragmented the dynamical regimes were over parameters, we used total variation: the sum of absolute differences between neighbouring elements (Methods). A larger total variation suggests sensitivity to parameter changes, and a lower total variation indicates greater robustness and stability to parameter variation. We found that the larval CoNNs produced smoother MLE dynamical landscapes than random networks (Figure 4C left, *p* = 0.004). Similarly, the total variation of Lyapunov dimension for CoNNs was lower than random networks (Figure 4C right, adult *p* < 0.001).

However, the increased dynamical robustness and stability across varying parameters may come at the cost of lower dimensional dynamics: we found that the Lyapunov dimension was consistently lower in CoNNs than random networks (Figure 4A), suggesting that the connectome intrinsic dynamics evolve on more restricted, low-dimensional subspaces. Further, the Lyapunov dimension was lower for the sparser larva-based networks than adult ones, in both connectome and random cases.

To test if the low Lyapunov dimension of the CoNN dynamics translated into other measures of dynamical dimensionality, we performed principal component analysis (PCA) on the network activity and calculated mean neuronal correlation. The cumulative variance explained (Figure 4D) revealed that dynamics in the CoNNs were captured by a smaller number of principal components compared to random comparison networks. We calculated the significance of this comparison by computing the participation ratio (PR) of the proportion of variance explained over the top 15 principal components for each network. The higher the PR value is, the more distributed the quantity is. We found that the CoNNs have significantly lower mean PR values (2.77 and 4.65 for larva and adult) than the random networks (5.43 and 6.87 for larva and adult) with *p*-values of 0.004 and *p* < 0.001 for larva and adult comparisons, respectively, indicating that the dynamics are more distributed and higher-dimensional in the random case. The larva CoNNs exhibited a significantly lower mean PR than the adult ones (*p* = 0.002, Mann-Whitney U test).

Similarly, when we computed the mean pairwise correlations across all neurons during the above dynamics, we found that — in keeping with the lower-dimensional dynamics — the mean correlation was higher for larval CoNNs than random comparison networks (Figure 4E), with *p* = 0.004 and *p* < 0.001 for the larva and adult cases, respectively.

The Configuration Model network dynamics displayed a mean dimensionality between that of the connectome and Erdős-Rényi networks (Figure S25F and G).

### Eigenspectral Properties

The eigenspectrum of a network plays a key role in shaping dynamics (62, 30, 63), and particularly in sparse networks (64). The full baseline eigenspectra of the CoNNs differed markedly from those of equivalent random networks. Across CoNNs, the eigenvalues clustered more tightly around zero — both in their real and imaginary components — whereas random networks display a broader, more uniform spread (Figure S24A). Larger values of real parts correspond to longer temporal transients and hence longer timescales of integration. Similarly, larger values on the imaginary axis support richer frequency representations in neural activity (29, 65). Thus, the relatively narrow eigenspectra of CoNNs may constrain their ability to retain information over long timescales or to represent diverse temporal features.

Example eigenspectra from a connectome and random network are shown in Figure S24B. To further illustrate the functional impact of these differences and exemplify the impact of the eigenspectrum on the timescale and diversity of transients, we provided an identical initial pulse to both network types and allowed the activity to decay under intrinsic dynamics. Figure S24C demonstrates that the random network exhibit longer-lasting transients and greater frequency diversity, consistent with its broader eigenvalue spread. This comparison highlights how the structural constraints encoded in networks shape the richness and timescales of their responses.

## Discussion

We used the Echo State Network framework to investigate how the connectivity of larval and adult *Drosophila melanogaster* connectomes shapes network dynamics and computation. Across a suite of tasks, connectome-based ESNs (CoNNs) achieved performance comparable to conventional networks, while exhibiting distinct advantages in efficiency and robustness.

Our work has expanded existing results on biologically-inspired ESN performance (25, 26, 34, 37, 33, 66, 67) to go beyond the typical tasks of memory and chaotic time-series prediction, incorporating a larger range of computational tasks including paired variants of working memory, decision-making, non-chaotic time-series prediction, and chaotic time-series prediction.

### Efficiency

CoNNs demonstrated consistently greater efficiency. When the networks were constructed with equivalent spectral radii, the CoNNs showed smaller synaptic weights and a lower total “wiring cost” than random networks. When accounting for this synaptic weight cost, CoNNs outperformed random networks on most tasks, indicating a possible favourable trade-off between wiring cost and functional capacity (33). This aligns with the idea that biological networks are organised to optimise both computation and resources.

This efficiency is further reflected in the reduced total neural activity variance across tasks, suggesting possible lower energetic demands. CoNNs also showed more specialised, narrowly distributed neural engagement: weighted task variance revealed that neurons in CoNNs were more specialised, with lower participation ratios both across tasks per neuron and across neurons per task. In contrast, random networks distributed their activity more broadly. Thus, the connectome topology supports sparse, task-specific coding (68–70), where computation is concentrated in smaller subsets of neurons, while maintaining performance. The relationship between local network features such as node degree, clustering, and self-recurrency with neural engagement suggests that these structural properties contribute to task-relevant activity. While these correlations are present in both connectome and random networks (uncovering important general principles of structure and function in specific tasks), they are stronger in CoNNs, where such features are markedly more prevalent, indicating their functional role in promoting efficient, selective engagement. The importance of local network features in ESN computation is supported by other findings (30, 29, 71, 72, 45).

The fact that particular annotated biological cell types correlated with weighted task variance may simply point to the link between cell type and the structural features identified of self-recurrency, clustering, and node degrees (43, 44); however this in itself may further indicate how structural network features are relevant for biological network dynamics and functions.

### Robustness

The selective neural engagement of CoNNs contributes directly to enhanced robustness. Under structured pruning of neurons, CoNNs preserved task performance, spectral radius, and Lyapunov exponent more effectively than random networks. The narrower concentration of task-relevant contribution on smaller groups of neurons appears to buffer the networks against structural perturbations, linking weighted task variance and local network features with robustness and resilience.

Robustness is also evident in the dynamical regimes of CoNNs under parameter perturbation. Larva CoNNs remained closer to the edge of criticality under variation of input and recurrent weight scaling, showing smoother dynamical landscapes. This results in more stable and predictable dynamics (28, 35, 38, 64). The enhanced self-recurrency in larval networks explained this increased resilience.

However, this robustness and efficiency comes at a cost: CoNNs have a reduced dimensionality and a reduced dynamical expressiveness. This is consistent with biological findings which show that, even during complex behaviours, neural population activity typically occupies a low-dimensional subspace compared to the high-dimensional space of recorded neural activity (73– 75). This limitation is evident in the network dynamics and eigenspectra, and diminished performance on working memory tasks. This highlights a functional limitation on internal representations and memory (76) in CoNNs. This suggests that there is a trade-off between high-dimensionality, stability and robustness and that the connectome topology favours one type of task over another. Biological networks appear tuned to operate in a regime of efficient, low-dimensional, and resilient dynamics.

### Developmental comparisons

Larval networks displayed a greater degree of robustness to structural and parameter perturbations than adult networks. This potentially reflects the constraints and demands of early developmental stages, where neural circuits may prioritise coarse-grained, robust organisation to support reliable wiring, before transitioning towards more finely-tuned regimes in later development. Indeed, larval and adult connectomes exhibit distinct anatomical organisation, implying a functional network transition across development (43, 77).

### Limitations and Future Directions

There are several limitations in the methodology and interpretations of this research.

We used the adult *hemibrain* connectome (as opposed to the full adult brain) as it is one of the most widely-used connectome datasets of manageable size. However, the hemibrain represents only a fraction of the full adult brain, primarily covering one hemisphere of the central brain and excluding certain peripheral regions. Consequently, our results will not capture network features associated with bilateral or sensory-motor integration present in the complete system.

We used a hierarchical stochastic block model decomposition of the connectomes to obtain smaller, biologically meaningful subnetworks for analysis. This approach ensured that our subdivisions reflected structure in the *Drosophila* connectome (e.g. 41), rather than arbitrary partitions. We selected an intermediate level in the hierarchy to balance manageable network sizes with a retention of internal substructure. While this approach is data-driven and aims to understand general principles of computation in sparse networks (rather than specific *Drosophila* network functions), the resulting subnetworks are model-dependent and may not correspond to subnetworks employed in the real brain.

Relatedly, a potential limitation of our approach is over-interpreting the biological realism or relevance of CoNNs. By abstracting from biophysical properties to a weighted adjacency representation, where synaptic weights are drawn from a distribution, and neuron dynamics are modelled with a simple tanh nonlinearity, we omit details of synaptic dynamics, neuromodulation, and plasticity mechanisms that may critically shape real neural computations. Our approach thus reflects structural potential rather than full physiological function. Further, connectivity alone is not sufficient to uniquely determine network dynamics and function (78). Many different parameterisations of a recurrent network with the same structural wiring can yield distinct dynamical regimes and functional outcomes. Our CoNNs assume that the fixed wiring of the larval or adult connectome directly constrains meaningful dynamics in a biological way, yet results suggest that additional factors — such as neuron-specific gains, biases, and synaptic weights shaped by experience — may be essential to fully capture the biological system’s behaviour (78, 79). Thus, while the connectome provides an anatomically grounded scaffold, our findings should be interpreted as characterising one possible configuration of structure, dynamics, and computational function consistent with its topology, rather than the unique or “true” biologically realised one.

These limitations naturally suggest various directions for future work. Crucially, integrating more biologically-informed detail (such as weight distributions, synaptic plasticity rules, or neuron-type-specific dynamics) in a controlled manner would allow us to disentangle their relative contributions to brain computation, and could help link connectomic organisation more directly to function (80, 81). Ultimately, combining connectome topology with biologically grounded dynamics would offer a promising route toward unifying the threads in this research of network neuroscience, recurrent computation, and realistic biological function.

## Methods

### Echo State Network Setup

We use the Echo State Network (15) formulation of reservoir computing throughout this work. The ESN reservoir computing setup consists of an input layer, a fixed recurrent reservoir, and an output layer (Figure 1). The ESN framework is defined by the following:

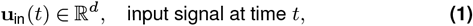

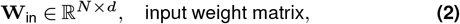

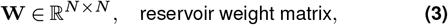

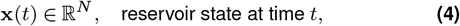

where *d* is the dimension of the input signal, *N* is the number of network nodes, and **W**_in_ and **W** are initialised with weights uniformly sampled from [− 1, 1]. The input layer **W**_in_ is scaled by a hyperparameter *ϵ*, the optimal value of which may vary according to task. The network states **x** evolve as:

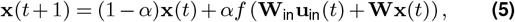

where *α* is the leak term, and the non-linear activation function *f* adopted in this work is tanh(·), the hyperbolic tangent. The output layer maps the network state to the desired output using a readout weight matrix **W**_out_ ∈ ℝ^*k×N*^ :

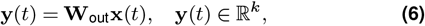

where *k* is the dimension of the output signal. Ridge regression is used to train the output weights **W**_out_. We define the matrix of network state time series **X** and the target outputs **Y**\*:

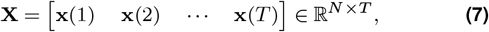

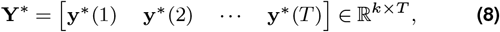

where **y**\*(*t*) is the target output at time *t*, and *T* is the number of time steps. The readout weights are computed using Ridge Regression as:

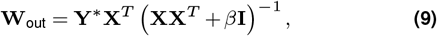

where *β >* 0 is the Tikhonov parameter (a regularisation parameter), and **I** is the identity matrix.

To control the dynamics of the reservoir during tasks and various analyses, we normalised the weight matrix **W** to have a specified spectral radius *ρ* = 0.99 (unless otherwise specified). This is a common spectral radius choice in ESN literature, and is further discussed in the section on Parameters. The spectral radius of **W**, denoted as *ρ*(**W**), is the largest absolute eigenvalue of **W**. To set a desired spectral radius *ρ*_target_, we first compute *ρ*(**W**) and then rescale **W** as

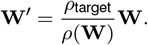

### Connectome Echo State Network Generation

We used the synapse-resolution larval (21) and adult *Drosophila melanogaster* (22) connectome to construct connectome-based ESNs (Figure 1 and S1). The *Drosophila* connectomes range from approximately 3,000 to over 20,000 neurons in the larva and adult hemibrain respectively, and therefore we constructed smaller subnetworks for use as ESNs. The *Drosophila* neural network exhibits a hierarchical modular structure according to type, class and function (41). In light of this, we therefore built connectome-based ESNs from subnetworks that we discovered in the *Drosophila* connectomes via a hierarchical stochastic block model (42, 26, 25). The hierarchical stochastic block model (HSBM) is a generative probabilistic model that recursively partitions a network into blocks (or communities) by maximising the posterior probability of the model given the observed adjacency matrix. It assumes that connections between nodes are governed by block memberships, and infers the most likely hierarchical structure using Bayesian model selection. We implemented the HSBM using the graph-tool Python package (42), which efficiently infers the hierarchy and allows extraction of subgraphs corresponding to distinct functional modules.

We chose subnetworks in hierarchical layers which gave appropriately sized communities (between 100–400 neurons) for our reservoir computing purposes, while maintaining sufficient subnetwork structure. In line with other research on biological reservoir networks (e.g. (34)), we randomised the weights of the connectome subnetworks (sampled uniformly from [−1, 1]) while preserving the topology (i.e. who connects with whom). This provided us with 9 larval *Drosophila* ESNs and 28 adult *Drosophila* hemibrain ESNs. We generated 30 instantiations with randomisation of weights for each of these connectome ESNs, preserving the topologies (Figure S1).

### Null Model Echo State Network Generation

For baseline comparison we used two random network models, namely, the conventional Erdős-Rényi random graphs and Configuration Model networks (82, 83). The Erdős-Rényi networks provide a reservoir network standard (18, 24, 84), while the Configuration Model provides a comparison where the degree distribution is matched to connectome networks, but is otherwise random. The Erdős-Rényi model generates a random directed graph by connecting each pair of nodes with independent probability *p*, where *p* is chosen to yield a desired sparsity. The node degrees of Erdős-Rényi networks follow binomial distributions.

Both Erdős-Rényi and Configuration Model networks were initialised with equivalent sparsities and spectral radius to the connectome networks. We generated 30 instantiations of ESNs for all of the networks.

### Statistical Significance

Unless otherwise stated, the statistical significance of group comparisons was assessed using the paired non-parametric Wilcoxon signed-rank test. We adopt a significance level of *α* = 0.05.

### Tasks and Performance Measures

Using the reservoir computing setup described, we can train ESNs to perform a variety of tasks. For this study, we selected four broad categories of tasks, namely: working memory, decision-making, timeseries prediction, and chaotic time-series prediction. For each category, we implemented two distinct task types, resulting in a set of eight tasks altogether (Table 1):

**Table 1.** Eight tasks grouped in their categories of memory, decision-making, time-series prediction, and chaotic time-series prediction.

|  |
| --- |
| (1a) Memory Capacity |
| (1b) Sequence Recall |
| (2a) Two Input Decision-Making |
| (2b) Delay Decision-Making |
| (3a) Trigonometric Series Prediction |
| (3b) Lotka-Volterra Prediction |
| (4a) Lorenz Time-Series Prediction |
| (4b) Rössler Time-Series Prediction |

These semi-cognitive-inspired tasks were selected to encompass a wide range of practically and conceptually different factors, each placing different kinds of demands on the reservoir framework. They were chosen with the purpose of being able to draw connections and contrasts across similar and distinct tasks, inferring possible relations between network structure and function. See Supplementary Information 2 for more detail on the task frameworks and performance measures.

### Wiring Cost-normalised Performance

Previous biologically-relevant reservoir computing research (e.g. (33)) has argued that the wiring cost of networks should be taken into consideration when comparing the performances of CoNNs and equivalent null models, such as Erdős-Rényi networks. They point out that the brain is a physically constrained and embedded network, with finite metabolic and material resources; brain networks have a prevalence of shorter, low-cost connections (3, 41). When account is taken of wiring cost, (33) demonstrate that the connectome networks outperform the null models across all dynamical regimes, suggesting that brain network topologies optimise the trade-off between computational capacity and cost. Inspired by this, we normalised the task performances of networks by the sum of the absolute values of their weights.

### Energy use during tasks

To quantify activity-dependent computational cost, we computed for each task the total squared network activity, 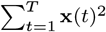, where **x**(*t*) denotes the network state at time *t*, and *T* is the length of neural activity. This measure captures the cumulative magnitude of neural activation during task execution and serves as a proxy for activity-based energy expenditure. It effectively measures how much overall the state deviates from inactivity.

### Parameters

The typical hyperparameters of interest in reservoir networks are reservoir size *N*, spectral radius *ρ*, sparsity *s*, input scaling factor *ϵ*, leakage rate *α*, and regularisation coefficient *β* (Tikhonov parameter). The sizes and sparsities of the networks were determined by the connectome, and therefore needed not be optimised. Based on the literature (18, 33, 19, 66, 39, 85) and our experience, we set the spectral radius to be close to (but strictly less than) unity (i.e. *ρ* = 0.99) which effectively sets the dynamics of the network close to criticality. We ran a grid search on the remaining parameters (*ϵ* ∈ [0.01, 0.02, 0.03, 0.04, 0.05, 0.1, 0.2, 0.3, 0.4, 0.5], *α* ∈ [0.1, 0.2, 0.3, 0.4, 0.5, 0.6, 0.7, 0.8, 0.9, 1.0], and *β* ∈, [10^−2^ 10^−3^, 10^−4^, 10^−5^, 10^−6^, 10^−7^, 10^−8^]) for all ESNs and across all eight tasks outlined above. The optimal set of parameters for a given task were used throughout this study.

### Weighted Task Variance

Task Variance is a measure that has been used to analyse the contribution and engagement of a neuron during an activity. It has been applied to determine whether individual units in a network are selective to different tasks, or whether units tended to be similarly selective to all tasks (49). The task variance of unit *i* on task *A* is defined as

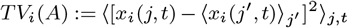

where *x*_*i*_(*j, t*) is the activity of node *i* on time *t* of trial *j*. Essentially, to compute task variance for a node *i* in Task *A*, one runs the network for many task iterations and conditions, and then calculates the variance across trials at each time point for a specific unit, and then averages the variance across all time points to get the final task variance for given unit *i*.

We introduce an augmented measure, called the *Weighted* Task Variance (WTV), where the weighted task variance of unit *i* on task *A* is defined as:

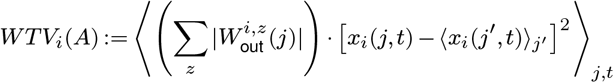

where 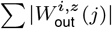 is the weighted contribution of the trained output layer in trial *j*, across output nodes *z* to given network node *i*. In this formulation, the importance of a given node’s task engagement is augmented by the size of its contribution to the final output. We calculated the weighted task variance of each node in all connectome and random networks, across the six different tasks.

### Participation Ratio

In order to be able to compare the distribution and level of neural engagement in the weighted task variance measures, we used the participation ratio (86, 87), defined as

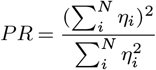

where *η*_*i*_ are the weighted task variances of neurons on a given task. Non-normalised participation ratio (PR) ranges from 1 (all centred on one mode) to *N* (uniformly spread across all modes). We used the participation ratio of weighted task variance (WTV) in two ways: Given an *T* × *N* weighted task variance matrix (where *T* denotes the number of tasks, i.e. 8, and *N* denotes the number of neurons in the network), we worked out the PR both across (i) columns and across (ii) rows (Figure 2A).

(i) That is, for every neuron in the network, we calculated the mean PR of WTV across all 8 tasks, which indicated how that neuron’s WTV was distributed. A higher PR indicates that the neuron has a more distributed WTV across the tasks, while a lower PR indicates that the neuron has a more narrow distribution across the tasks. This enabled us to determine which neurons were more task-specialised (lower PR) and those which were more task-generalised (higher PR).

(ii) Similarly, for each task, we calculated the mean PR of WTV across all neurons in the network. This enabled a comparison of how concentrated or distributed the neural contributions are *per task*, and indicated how sparse (lower PR) or distributed (higher PR) the neural contribution was on each task.

### ystematic Pruning of Nodes from Networks

Using the weighted task variance measures, we were able to define an ordering on the nodes within each network for a specific task. This ordering captures the relative importance of a neuron’s contribution for that task. We conducted a systematic lesioning process, pruning neurons beginning with the lowest ranked to the highest ranked, until the entire network is eliminated. As we iteratively removed nodes from the network, we measured task performance to indicate how pruning affects the capacity of the network. This enabled us to verify that the weighted task variance captures a measure of the importance of a node’s contribution for a task. We also measured how close the network states remain to “the edge of chaos”, or the degree of criticality in the network dynamics, captured by the maximum Lyapunov exponent. This provides an indication for how the structures of the connectome and random networks either successfully or unsuccessfully manage to sustain criticality in their activity, and how each node contributes to this functionality.

For comparison, we also pruned neurons from the networks randomly, instead of systematically in order of weighted task variance.

### Linking Network and Neuron Features to Function

Using the weighted task variance measure, we quantified how individual neurons contribute to distinct computational tasks. To interpret these contributions, we analysed how WTV correlates with several structural and biological features of neurons in the network:

- Node degree
- Self-recurrency
- Local clustering
- Biological cell-type annotation.

The **degree** of a node *i* is the number of edges connected to it. The indegree 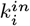 captures the number of incoming connections, while the out-degree 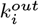 measures the number of outgoing connections. In and out node degrees are crucial for the integration and propagation of information across computational networks.

**Recurrency** refers to the tendency of a node to be part of directed loops that return to itself. These loops can be self-recursive (i.e., a node connects back to itself) or consist of multi-step cycles, such as two-step, three-step, four-step, and five-step loops that eventually return to the node. Recurrency is important in the context of recurrent neural networks as it influences the persistence and propagation of information within the network. We limited our results to self-recurrency, however, various analyses of multi-step recurrent loops showed similar qualitative results.

The **local clustering coefficient** (82) *C*_*i*_ quantifies how densely interconnected the neighbours of a given neuron *i* are. In the case of directed networks, *C*_*i*_ considers all possible directed triangles that include neuron *i*, accounting for the direction of edges in each triplet. Specifically, it measures the proportion of actual directed connections among neuron *i*’s neighbours relative to the number of all possible directed connections between them. A high local clustering coefficient indicates that the neighbours of neuron *i* are well connected among themselves, forming many directed loops, while a low value suggests a more sparse or feedforward-like local structure.

For these structural features, we computed the mean correlation of each measure with the WTV for all neurons. This gives us a picture of what network characteristics are most highly correlated with which tasks.

For various neurons within the connectome datasets, **cell-type** annotation labels were provided. We calculated the normalised mean WTV for each cell-type (controlling for the proportions of cell-types within a given network), also giving an indication as to what biological cell-type most strongly contributes to tasks.

### Characterising Dynamics of Networks

To explore the dynamical regimes of the ESNs, we computed various measures: the Lyapunov spectrum, the maximum Lyapunov exponent (MLE), the Lyapunov dimension, the proportion of variance explained, and the mean neural correlation.

### Lyapunov Spectrum and Maximum Lyapunov Exponent

The Lyapunov spectrum (54, 88) characterises the local stability properties of a dynamical system by quantifying how infinitesimal perturbations evolve along different directions in state space. For a system with state vector *X*(*t*) evolving under dynamics *X*(*t* + 1) = *f* (*X*(*t*)), the evolution of an infinitesimal perturbation *δX*(*t*) is governed by the system’s Jacobian *Jt* = *∂f/∂X*:

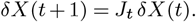

The long-term average exponential rates of divergence or contraction of these perturbations define the Lyapunov exponents 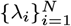, col-lectively known as the Lyapunov spectrum:

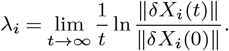

Positive exponents indicate directions of local expansion (chaotic dynamics), negative exponents correspond to contraction (stable directions), and zero exponents represent neutral stability. The overall shape of the spectrum therefore reflects the balance between order, memory retention, and sensitivity to perturbations in the system.

Among these, the largest exponent —-the *maximum Lyapunov exponent* —captures the dominant stability property of the dynamics. A positive value (*λ*_max_ *>* 0) signifies chaotic dynamics where perturbations grow exponentially, while a negative value (*λ*_max_ < 0) indicates asymptotic stability or convergence to fixed points or limit cycles. A system operating near *λ*_max_ ≈0 lies at the *edge of chaos* or *edge of stability*, a regime often associated with criticality —– balancing stability and flexibility to support rich, adaptive information processing. Criticality is a state of network dynamics which has been observed experimentally in neural recordings, where brain activity often hovers near a critical regime during cognitive tasks, and also computationally in artificial neural networks (58, 59, 61, 89), where networks tuned to the edge of chaos exhibit optimal performance in information processing, memory, and learning. As previously alluded to, criticality is a fundamental state of optimal network activity in ESNs (90, 33, 18, 91). For numerically calculating the Lyapunov spectra (Figures 4 and S19), we followed (56, 92) by iterating the autonomous network dynamics and propagating an orthonormal basis through the local Jacobian at each timestep. At each iteration, a QR decomposition was applied to the evolved perturbation vectors to maintain orthogonality, and the logarithms of the resulting stretch factors (diagonal entries of *R*) were accumulated over time. The time-averaged logarithmic growth rates yielded the full Lyapunov spectrum. The largest exponent from this spectrum corresponds to the MLE. From the ordered spectrum *λ*_1_ ≥*λ*_2_ ≥… ≥*λ*_*N*_, we also estimated the *Lyapunov dimension* (Kaplan–Yorke dimension) as

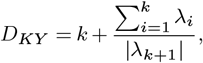

where *k* is the largest integer for which 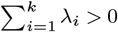. This provides an effective measure of the number of dynamically active degrees of freedom in the system.

These analyses allowed us to map the dynamical behaviour of ESNs in stability and dimensionality across architectures and parameter choices. We used total variation to capture how smooth or abrupt the transitions between dynamical regimes were. The total variation is computed as

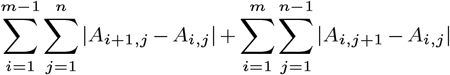

where *A* ∈ ℝ^*m×n*^ is the matrix of values across the *m* × *n* parameter sweep.

### Dimensionality of Input-driven Dynamics

To further quantify the dimensionality of the input-driven dynamics of the connectome and conventional ESNs, we drove each network with scalar noise input (uniformly sampled over [− 0.5, 0.5] with the input **W**_in_ layer scaled by *ϵ* = 0.01 over 1000 time steps) and analysed the resulting activity using principal component analysis (PCA). We recorded the network state matrix **X** ∈ ℝ^*N×T*^, where each column corresponds to the network state at a given timestep. We then applied PCA to **X** and extracted the leading 15 eigenvalues of the covariance matrix, which reflect the amount of variance captured along each principal axis. For each network, results were averaged across two independent simulations with different noise realisations. During these dynamics, we also computed the mean pairwise absolute correlation coefficient across all neurons.

## Code and Data Availability

Code and data relevant for this paper are available at https://github.com/jajmcallister/Conn_ESN_Paper/.

## Acknowledgements

J.M. is the recipient of a Northern Ireland Department for the Economy PhD Scholarship. We would like to thank Albert Cardona for useful comments.

## Supplementary Material

### Supplementary Note 1: Weight Distributions and Wiring Cost

The spectral radius *ρ* of a network is the largest absolute eigenvalue of its recurrent weight matrix, and it plays a significant role in the network dynamics. In random networks, the spectral radius controls the transition between contractive, stable dynamics and expansive or chaotic regimes, and is therefore a key parameter in ESN computational performance (8, 11, 12).

The spectral radius *ρ*(**W**) of a large non-symmetric random matrix **W** can be approximated using the Circular Law (10, 20, 21), which states that for an *N* ×*N* matrix with independent, zero-mean entries of variance *σ*^2^, the spectral radius is asymptotically approximated by 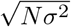.

Given that our weights are initially sampled **W**_*ij*_ ∼ 𝒰[−1, 1], we have that 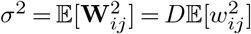, where *D* = 1 – *S* is the network *density* (for a given sparsity *S*). For a uniform distribution 𝒰[*a, b*], the second moment is 𝔼[*X*^2^] = (*b* −*a*)^2^*/*12, giving us that 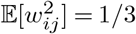, and thus *σ*^2^ = *D/*3 = (1 −*S*)*/*3. Therefore, the baseline spectral radius of our initially-constructed random networks (before rescaling *ρ*) can be approximated:

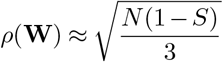

While the Circular Law from random matrix theory cannot be applied to the sparse and non-random topology of biological networks, (3) and (14) show that, for sparse random graphs, the spectral radius scales roughly as 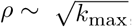, where *k*_max_ is the largest degree in the network. This means that the spectral radii of networks with hubs or heterogeneous connectivity (and therefore larger maximum degrees) can be expected to be larger than those of networks with relatively narrow degree distributions (and hence smaller maximum degrees) with equivalent weight distributions. Consequently, for our connectome networks — which have higher-degree nodes than the comparison random networks (Figure 1B) — one expects their spectral radii to be systematically larger, which is indeed what we find (Figure S1). This suggests that, even at identical sparsity and weight distribution, the intrinsic topology of the connectome achieves higher spectral radii relative to random networks.

**Figure S1.**
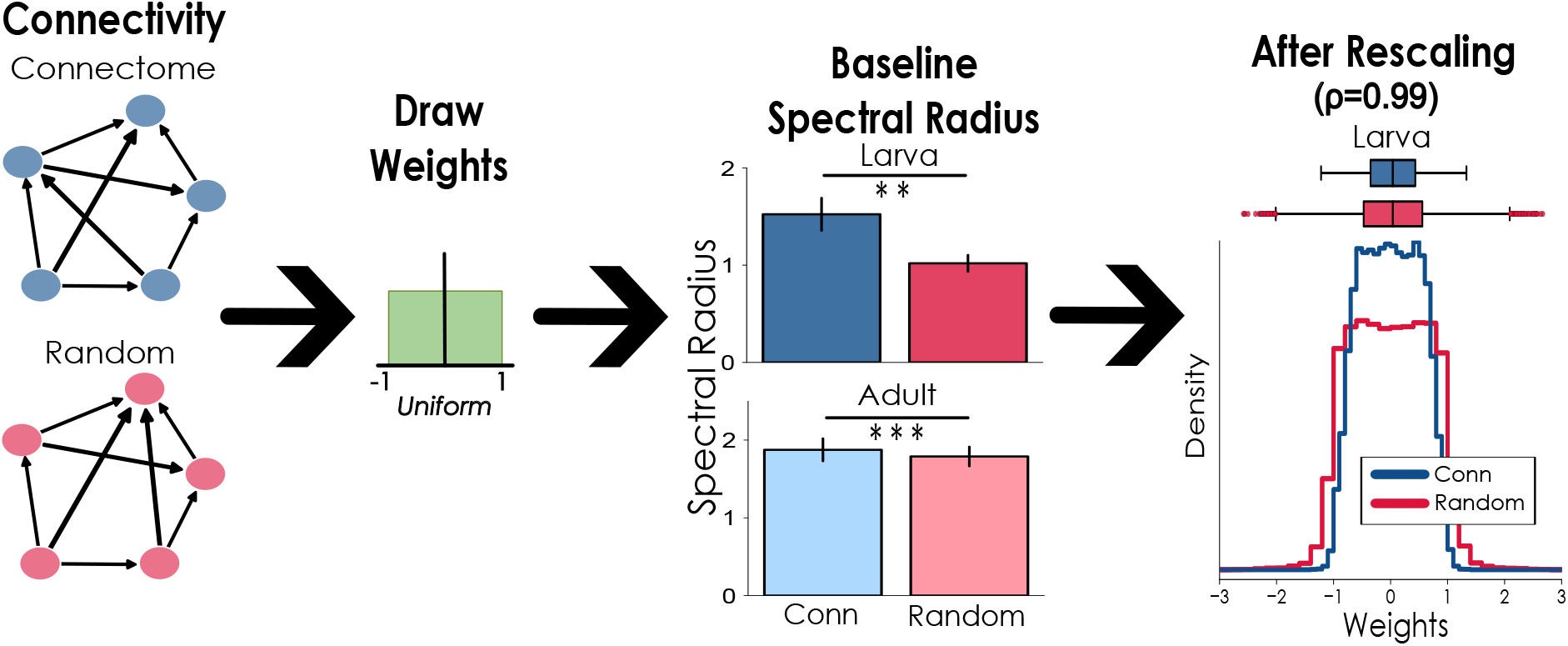
How connectome networks reduce wiring cost. The underlying network topology influences ‘baseline’ spectral radius, even when weights are drawn from the same initial distribution. When the ESNs are scaled to match a desired spectral radius *ρ* = 0.99, the resulting weight distribution for connectome ESNs is narrower than for random models: giving a lower ‘cost’.

Drawing weights from the same distribution for networks of different sizes, sparsities and topologies will produce varying network eigenspectra, and therefore different spectral radii. As mentioned, the spectral radius is a key parameter in ESN performance, and often is systematically controlled. For this reason, after the weights are uniformly drawn for the connectome and random ESNs (Figure S1), we scaled the weights to achieve equivalent spectral radii. Normalising the weight matrices means that, as a result of the larger baseline spectral radius of the connectome ESNs compared with the random null models, the resulting weights are smaller for the connectome networks than their random counterparts after rescaling (Figure S1).

A simple result is that the sum of the absolute values of the weight matrices — a basic proxy for total synaptic or wiring cost (1, 2) — is lower for connectome ESNs than random ones.

#### Supplementary Note 2: Task Details

##### 1a: Memory Capacity

This working memory capacity task is inspired by a plethora of other reservoir work (23, 6, 12, 8, 17). In this task paradigm, a random input sequence *X*(*t*) is presented to the network through a single input neuron. The network independently learns delayed versions of the input, producing multiple outputs. Each output *Y*_*τ*_ predicts the input *X*(*t*) delayed by *τ* time steps, i.e., *Y*_*τ*_ (*t*) = *X*(*t* − *τ*). The input values are randomly drawn from a uniform distribution, *X*(*t*) ∼ Uniform(−0.5, 0.5).

**Figure S2.**
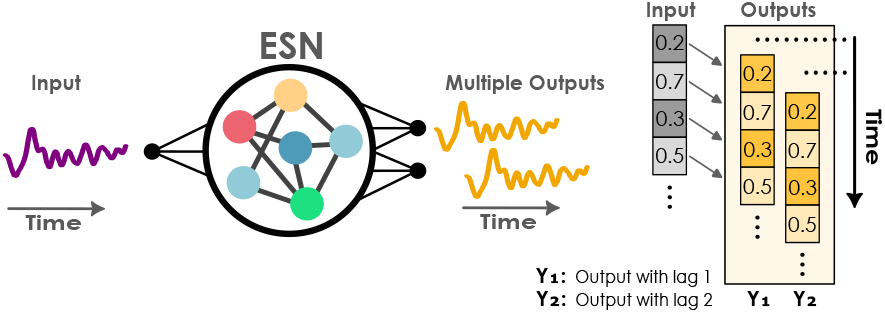
Illustration of the working memory capacity task.

**Figure S3.**
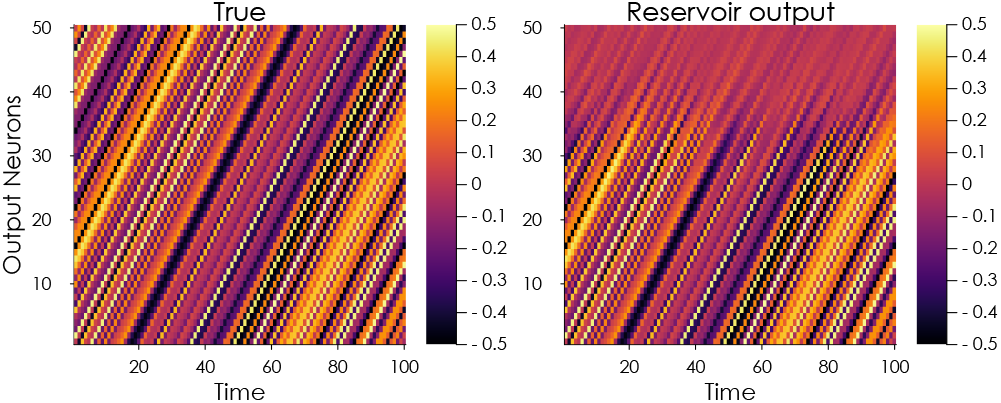
Working memory capacity task; the left indicates the true/target data, the right shows the network’s output. Notice the blurring or fading of the network memory performance.

The networks are trained for 4000 time steps and tested on the subsequent 1000. Each output is trained independently, and the performance, referred to as Working Memory Capacity, is calculated as the cumulative squared Pearson correlation coefficient (*ρ*) across all outputs:

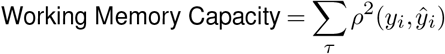

where *y*_*i*_ and *ŷ*_*i*_ denote the true and predicted values, respectively. Although such a simple input-output delay is a trivial task from an engineering point of view, we consider it a valuable benchmark task for reservoir networks, because any complicated task on time series data will need to be able to temporarily store information from the past in order to combine it with the present input.

**Figure S4.**
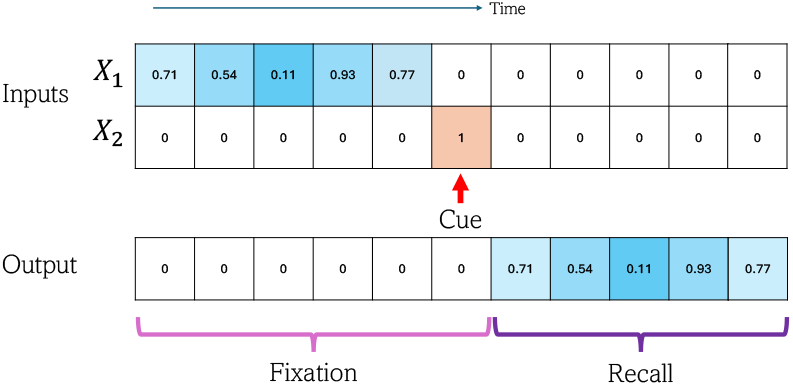
Sequence recall task

**Figure S5.**
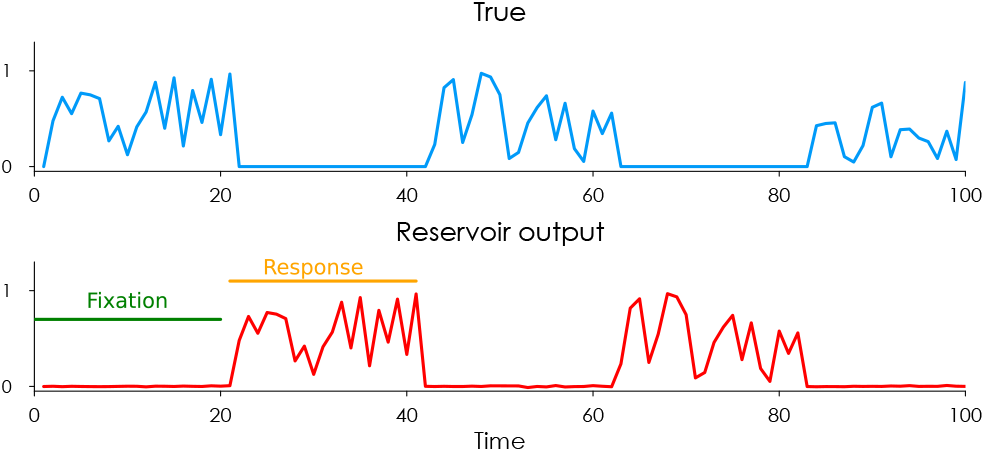
Example true data and actual network output for sequence recall task.

##### 1b: Sequence Recall

In the sequence recall task, the reservoir network is presented with two inputs: *X*_1_(*t*), a sequence of random numbers to memorise, and *X*_2_(*t*), a cue input. The cue input signals whether to fixate (output equal to zero) or to recall. During the recall phase, the network is required to output the memorised sequence corresponding to the *L* steps preceding the recall signal. The parameter *L* determines the pattern length and regulates the task’s difficulty. A single trial of the task consists of a fixation period followed by the recall period. The input values *X*_1_(*t*) are randomly drawn from a uniform distribution, specifically *X*_1_(*t*) ∼ Uniform(0, 1). Performance on a given trial is evaluated using the squared correlation between target and actual output.

In order to determine the sequence recall performance of a given network, we run the above task over increasing values of se-quence length *L* from 2 to 100. The sequence length just below which the squared correlation goes below a threshold (set at 0.6) gives the “sequence recall ability” of the network (**Figure** S6).

**Figure S6.**
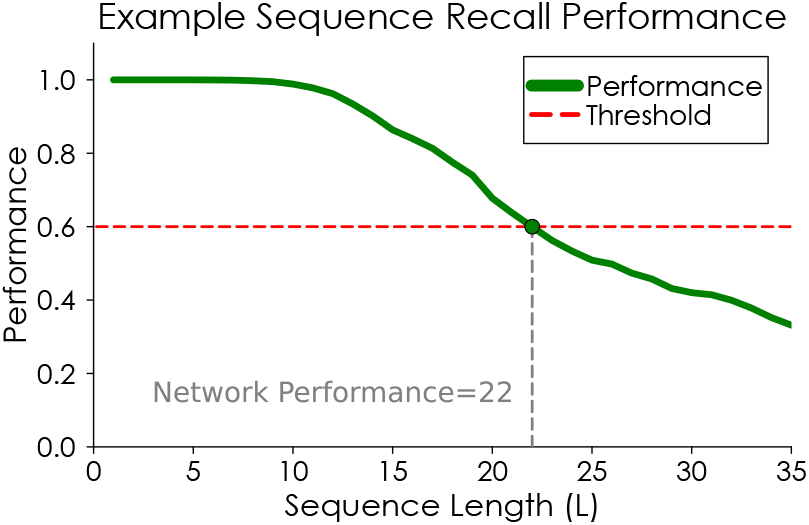
Illustration for calculating sequence recall performance for a network.

##### 2a: Two Input Decision-Making

In the two-input decision-making task, the network is presented with two input streams, *X*_1_(*t*) and *X*_2_(*t*), which consist of time-varying stochastic signals. The task requires the network to integrate the information from these two inputs over time and output a decision indicating which input has the higher average value. The output is represented by a single neuron, where a positive output (+1) indicates that *X*_1_(*t*) has a higher average, while a negative output (−1) indicates that *X*_2_(*t*) is higher. The input streams *X*_1_(*t*) and *X*_2_(*t*) are drawn from a Gaussian distribution, with a small bias added to one of them to create alternating dominance between the two signals. The size of this bias determines the difficulty of the task (how close or far apart the two signals will be). This task is inspired by classical perceptual decision-making tasks based on random-dot motion stimuli (18). In these tasks, a visual stimulus in the form of a set of moving dots is presented, with the motion of dots controlled by a coherence parameter. The task is to discern and discriminate the overall direction of motion among the dots, which may be either coherent (moving in the same direction) or incoherent (moving randomly). We instead use two input stimuli to represent momentary motion evidence for the two target directions (25, 22). Performance on a single trial is measured by decision accuracy, where the “decision” of the network is calculated by taking the signed mean of the output signal during a decision period.

**Figure S7.**
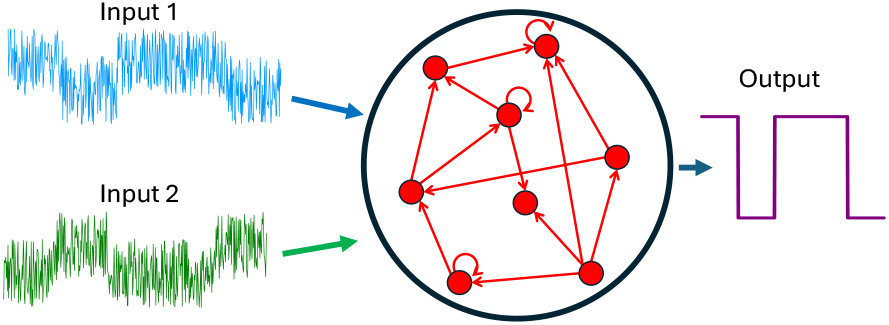
Illustration of the setup for the two input decision-making task.

**Figure S8.**
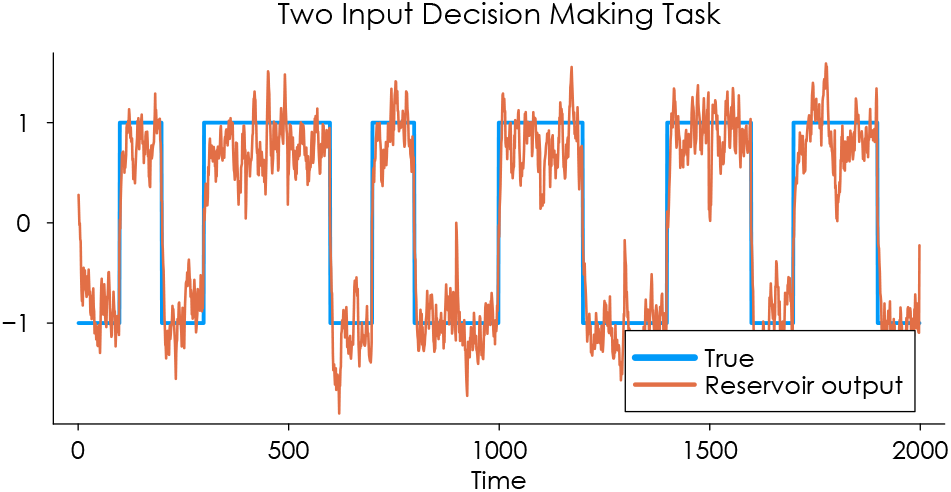
Example true data and actual reservoir output for two input decision-making task

To determine the performance of a given network on this task, we run the task with varying values of bias (between 0.01 and 1.0) and find the smallest level of bias (bias_min_) at which the network can accurately (within a threshold, set at 0.8) classify the inputs. We established a baseline bias at which all the networks could achieve near perfect accuracy (bias_baseline_ = 1.0); we therefore calculate the performance of a specific network to be bias_baseline_ −bias_min_.

##### 2b: Delay Decision-Making

The delay decision-making task alters the setup of the two input decision-making task by incorporating a stimulus and cue component (**Figure** S9), similar to the sequence recall task. In this task the network receives a continuous stream of noisy input via input neuron 1 over a fixed-duration stimulus period. The input signal fluctuates around zero with values drawn from a Gaussian distribution with mean +0.5 or −0.5 (which determines whether the stimulus is above or below 0 on average), and with a variance which determines the difficulty of the task. The network must integrate this information over time without producing an output during the stimulus phase (“fixate”). After the stimulus period ends, a cue signal is provided via input neuron 2, at which point the network must generate a binary decision: +1 if the time-averaged input was greater than zero and −1 if it was less than zero (“respond”). This task requires the network to maintain an internal representation of the accumulated evidence in a latent state and delay its response until prompted. Performance on a trial is measured by decision accuracy, where the “decision” of the network is calculated by taking the signed mean of the output signal during a given response period. This paradigm tests the network’s ability to perform working *both* memory-based integration and respond selectively based on an external cue.

To ascertain the performance of a given network on the delay decision-making task, we run the task with increasing values of variance (between 0.1 − 4.0). The variance (or “difficulty level”) just below which the accuracy drops below a threshold (set at 0.8) gives the performance of the network.

##### 3a and 3b: Trigonometric Oscillator and Lotka-Volterra Time-Series Prediction

We use a synthetic time series generated from a non-linear periodic function composed of multiple sinusoidal components:

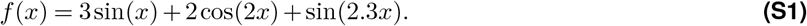

We sample the signal at a resolution of Δ*t* = 0.1, producing a long time series used for both training and testing. The first 2000 time steps are used for training, and a non-overlapping segment of 1000 steps are randomly selected for testing. For testing, the network is run in autonomous mode (**Figure** S11) after a “warm-up period” of 100 steps, which are driven by real data. Prediction quality is evaluated by the metric of valid prediction time — the time until which the normalised mean squared error remains below a threshold; here, set to 0.5. We calculate the normalised mean squared error (16),

**Figure S9.**
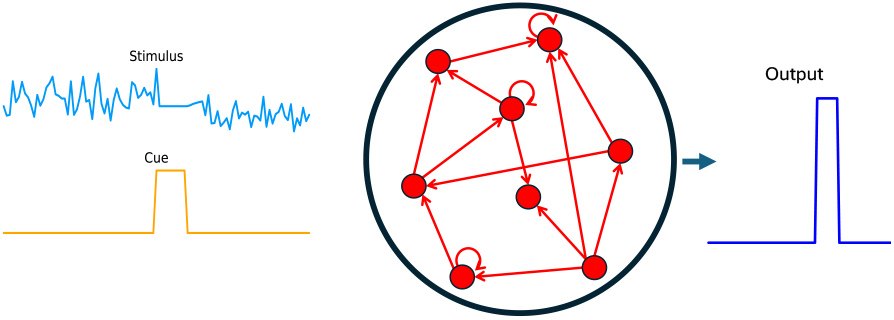
Illustration of the setup for the delay decision-making task.

**Figure S10.**
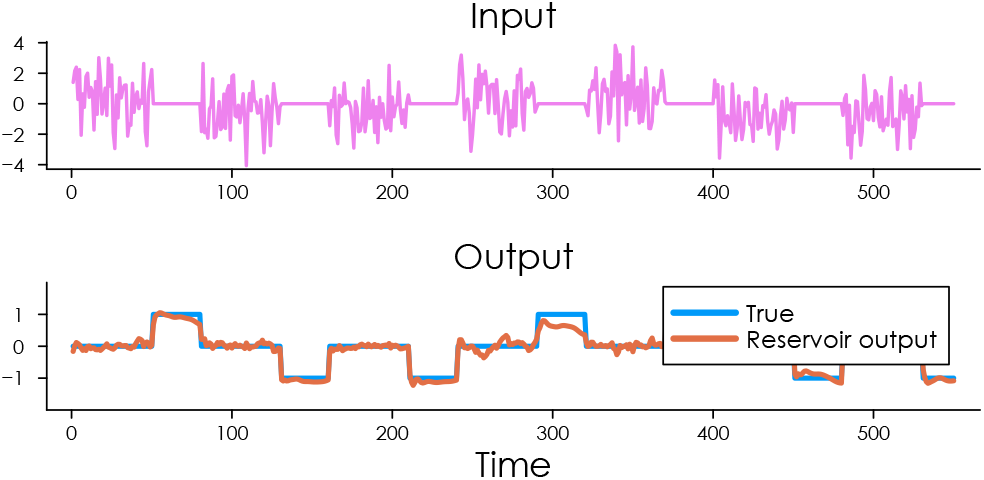
Example true data and actual network output for the delay decision-making task

**Figure S11.**
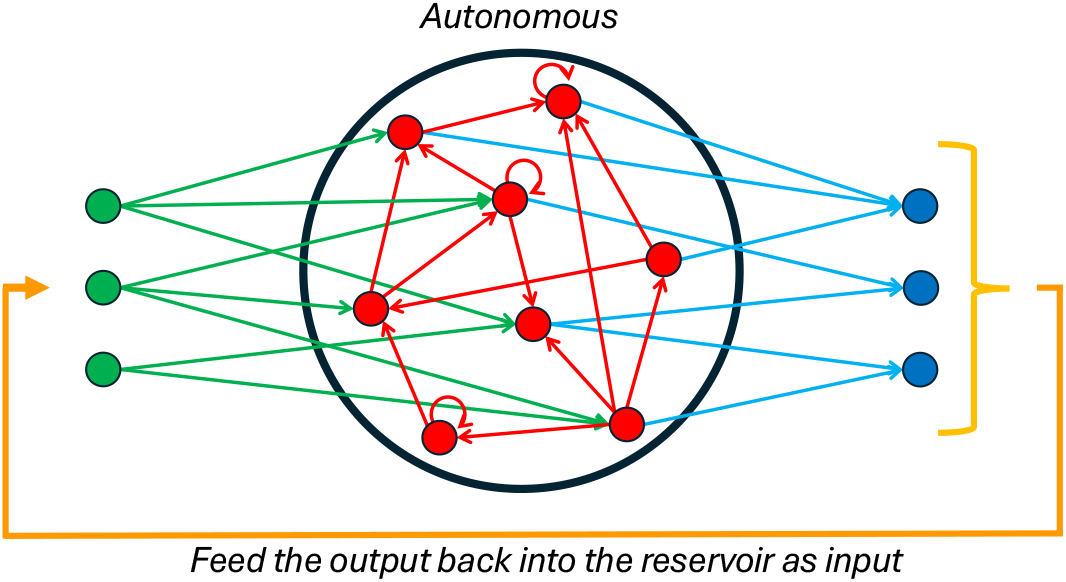
The autonomous setup for the reservoir network in time-series prediction tasks: The network output is fed back in as input.

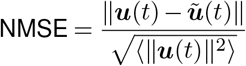

where *u*(*t*) is the true value and *ũ*(*t*) is the predicted value.

**Figure S12.**
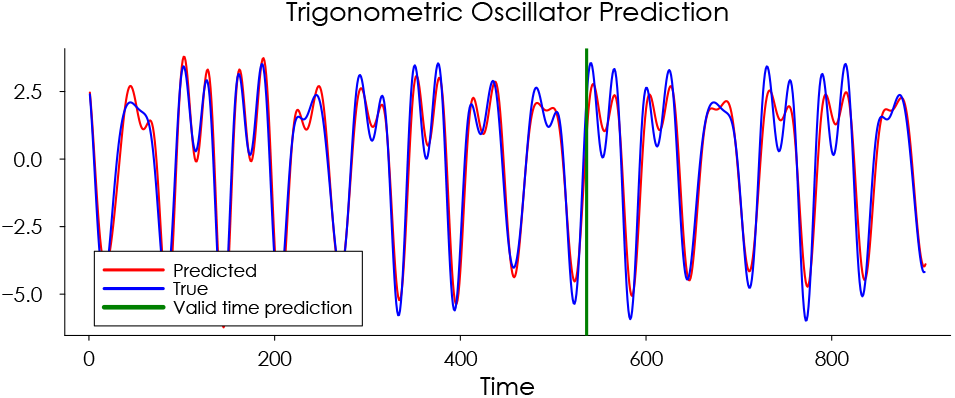
Example true data and actual network output for the trigonometric oscillator task

**Figure S13.**
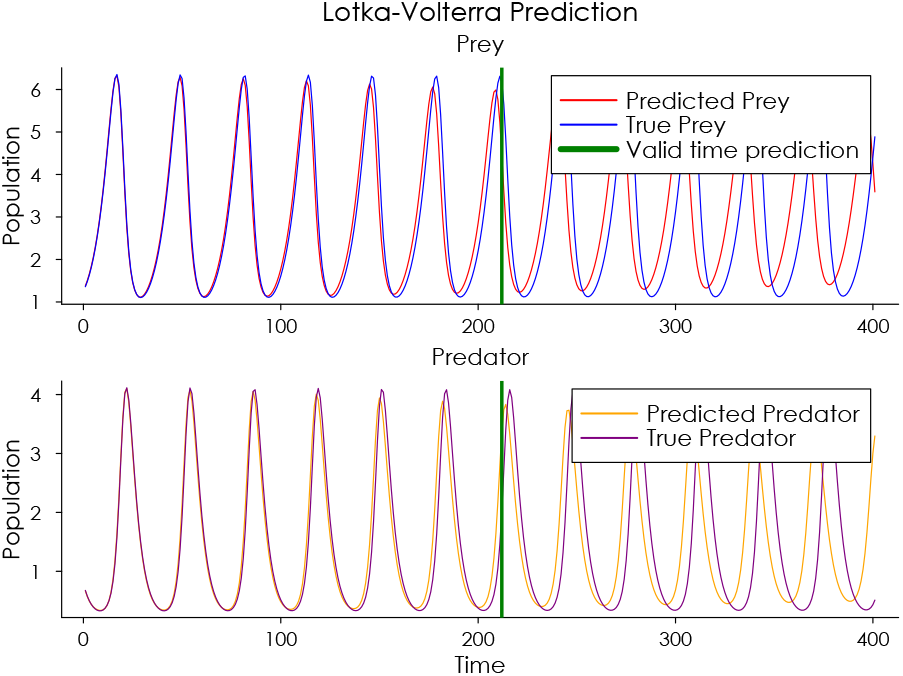
Example true data and actual network output for the trigonometric oscillator task

We use exactly the same approach with an alternative time-series task, the Lotka–Volterra system, which models predator–prey interactions through coupled non-linear differential equations. The system exhibits oscillatory dynamics governed by the equations

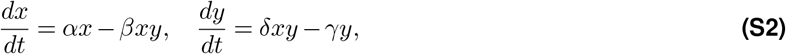

where *x*(*t*) and *y*(*t*) represent the prey and predator populations, respectively. The parameters *α, β, γ, δ* control the rates of growth, predation, predator mortality, and reproduction; these are set to 1.5, 1.0, 3.0, and 1.0 respectively. We simulate this system numerically to generate a two-dimensional time series.

##### 4a and 4b: Lorenz and Rössler Chaotic Time-Series Prediction

The canonical example of a reservoir computing task is prediction of the chaotic time-series data from the Lorenz system (16, 5, 4). The Lorenz system is notable for having chaotic solutions for certain parameter values and initial conditions. The system is described by the equations

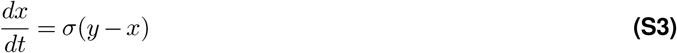

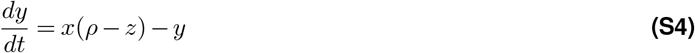

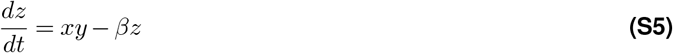

where *x, y*, and *z* are the state variables, and the parameters *σ, ρ*, and *β* are set to conventional values of 10, 28, and 8*/*3. We train the reservoir network on a set of Lorenz data, and then provide the system with a random point on the Lorenz trajectory, and measure the valid time (16) for which the reservoir network running autonomously (**Figure** S11) can accurately predict the true trajectory. We measure, in Lyapunov time, the valid time of the predicted trajectory, that is, the point at which the predicted trajectory exceeds a specified NMSE threshold, set at 0.5 in this study. We use the same approach with an alternative chaotic system, the Rössler system, described by the equations

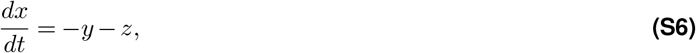

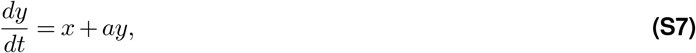

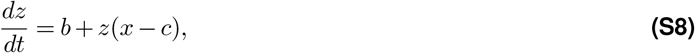

where *x, y*, and *z* are the state variables, and the parameters *a, b*, and *c*, which control the system’s behaviour, are set to conventional values of 0.15, 0.2, and 10.0 respectively.

**Figure S14.**
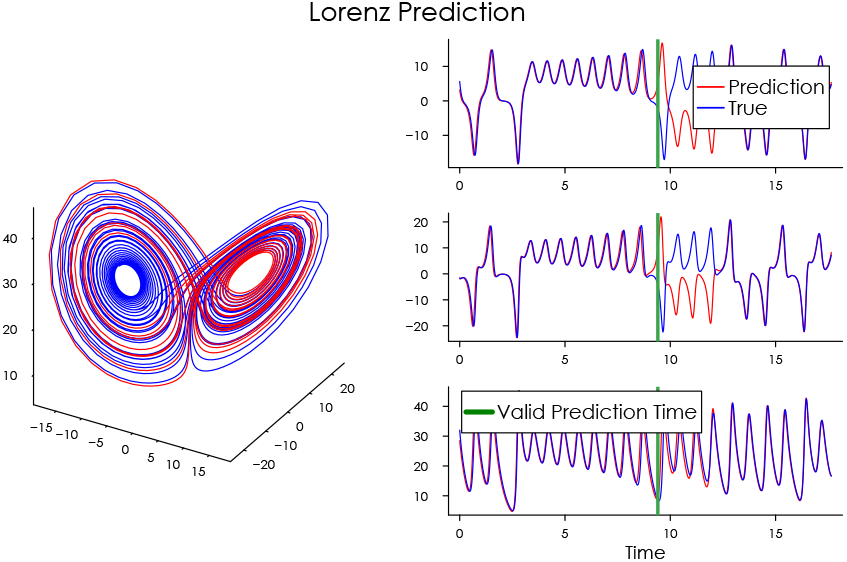
Illustration of the true and network prediction of the Lorenz system.

**Figure S15.**
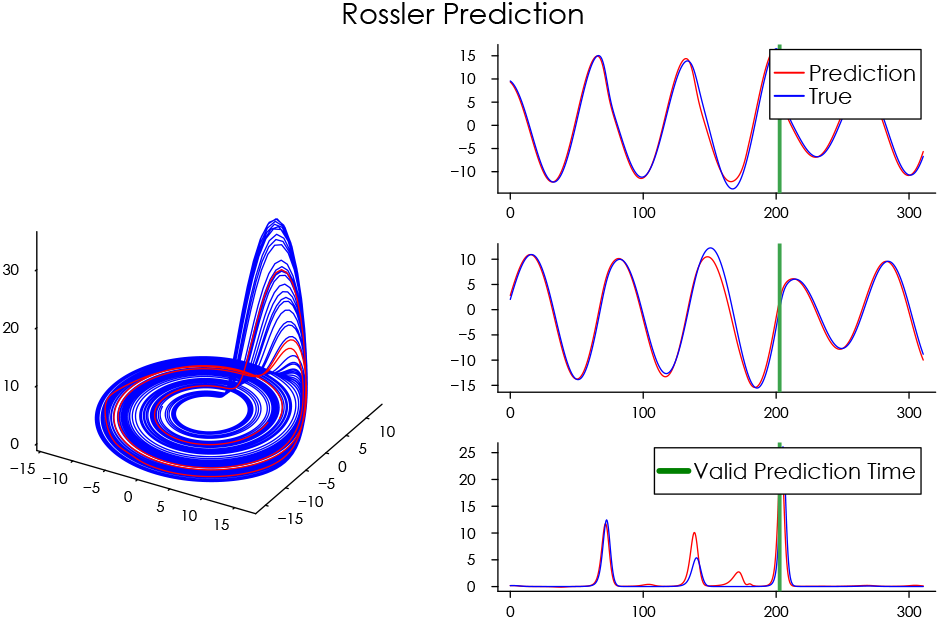
Illustration of the true and network prediction of the Rössler system.

#### Supplementary Note 3: Tasks – Performances, Energy Use, and Pruning

**Figure S16.**
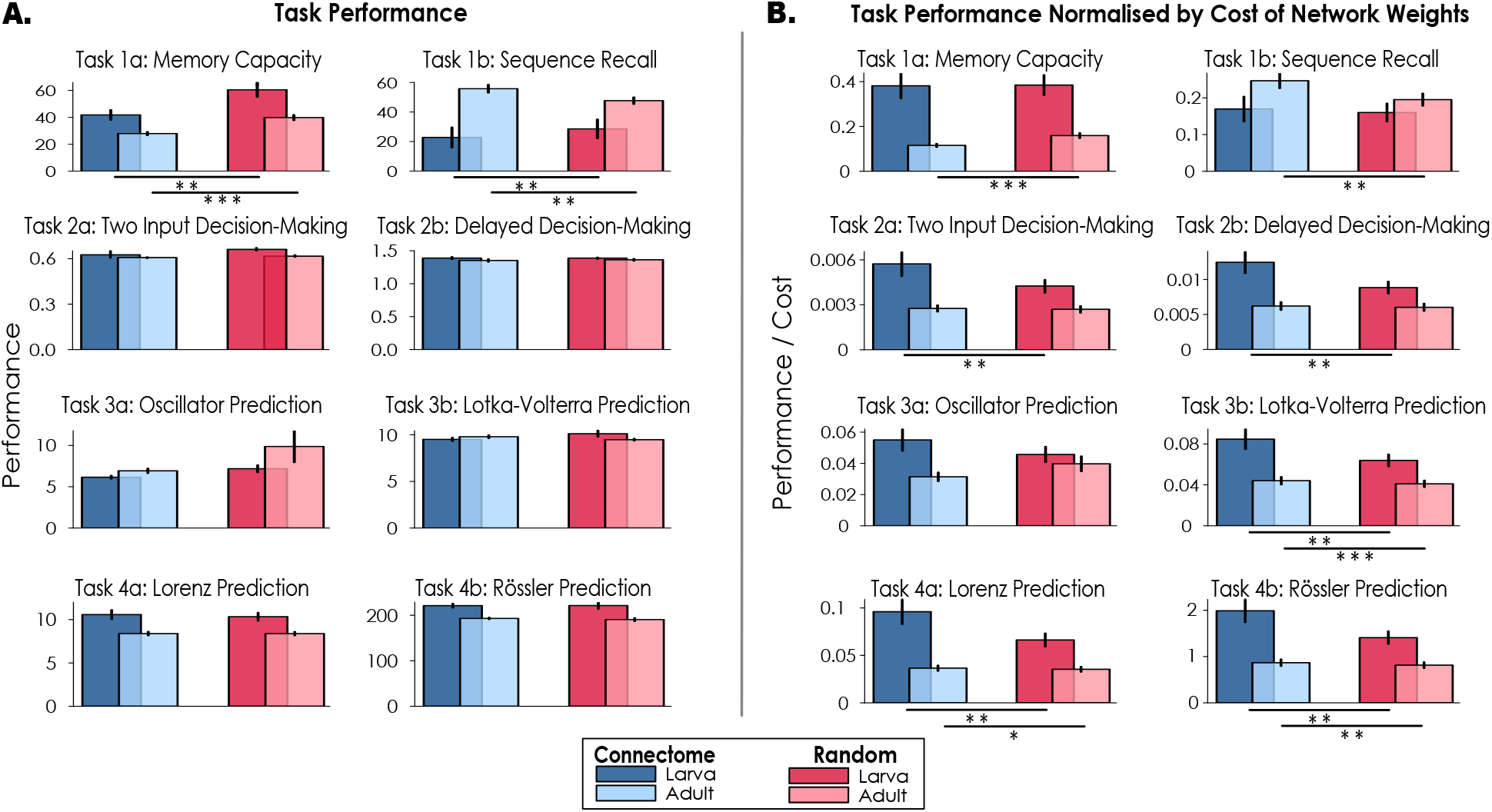
A Performances (and standard error of the mean) of connectome-based networks (CoNNs) and random networks on eight comp*u*tational tasks. Higher values mean better performance. Statistical significances found in the comparisons were *p* = 0.0039 and *p* = 7.45 *×* 10^−9^ (Task 1a, larva and adult), *p* = 0.0078 and *p* = 0.027 (Task 2, larva and adult). **B. Performances (and standard error of the mean) normalised by “wiring cost”**. Higher values mean better performance. Statistical significances found in the comparisons were *p* = 7.45×10^−9^ (Task 1a, adult), *p* = 0.0012 (Task 1b, adult), *p* = 0.0078 (Task 2a, larva), *p* = 0.0039 (Task 2b, larva), *p* = 0.0039 and *p* = 0.00079 (Task 3b, larva and adult), *p* = 0.0039 and 0.038 (Task 4a, larva and adult), and *p* = 0.0039 and *p* = 0.0027 (Task 4b, larva and adult). (* *p* < 0.05, * * *p* < 0.01, * * * *p* < 0.001)

**Figure S17.**
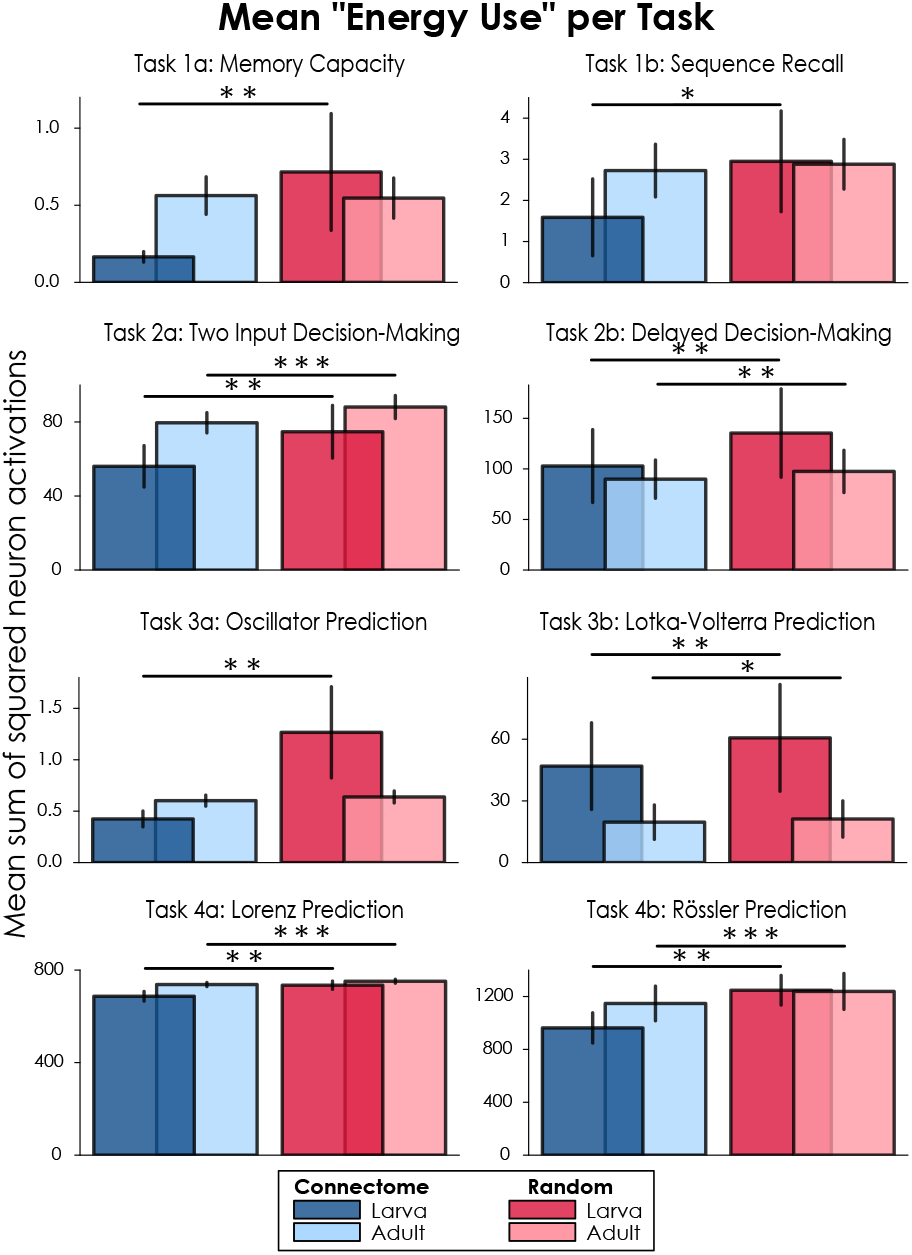
“Energy cost” (and standard error of the mean) per Task: Mean sum of squared neuron activations during the respective tasks. Lower values mean less overall neural activation, and hence lower “energy”.

**Figure S18.**
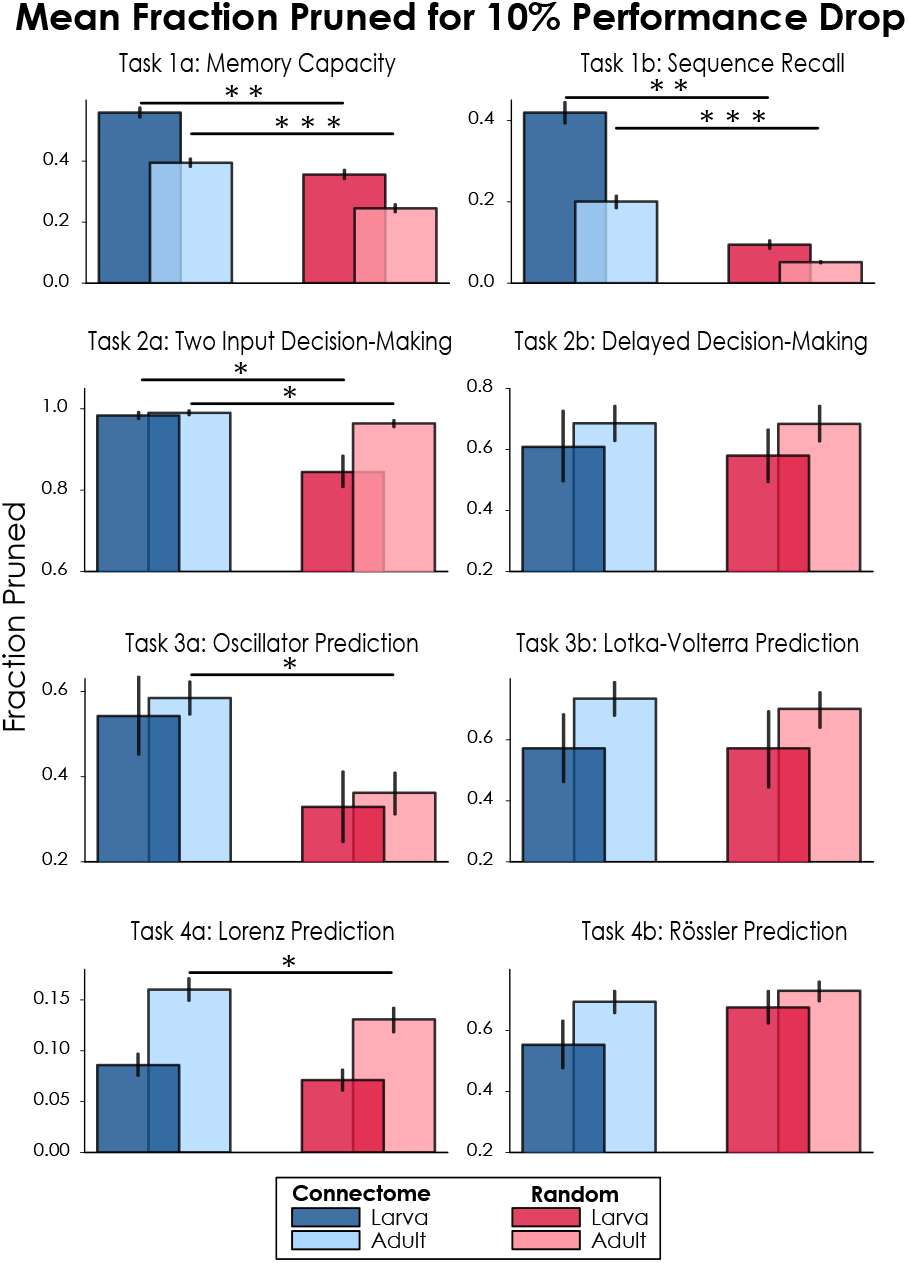
Mean fraction of networks removed (and standard error of the mean) before a 10% decrease in performance, as neurons are removed in order of increasing weighted task variance. Higher values mean the networks are more robust to pruning. (* *p* < 0.05, * * *p* < 0.01, * * * *p* < 0.001)

#### Supplementary Note 4: Maximum Lyapunov Exponent under Pruning

**Figure S19.**
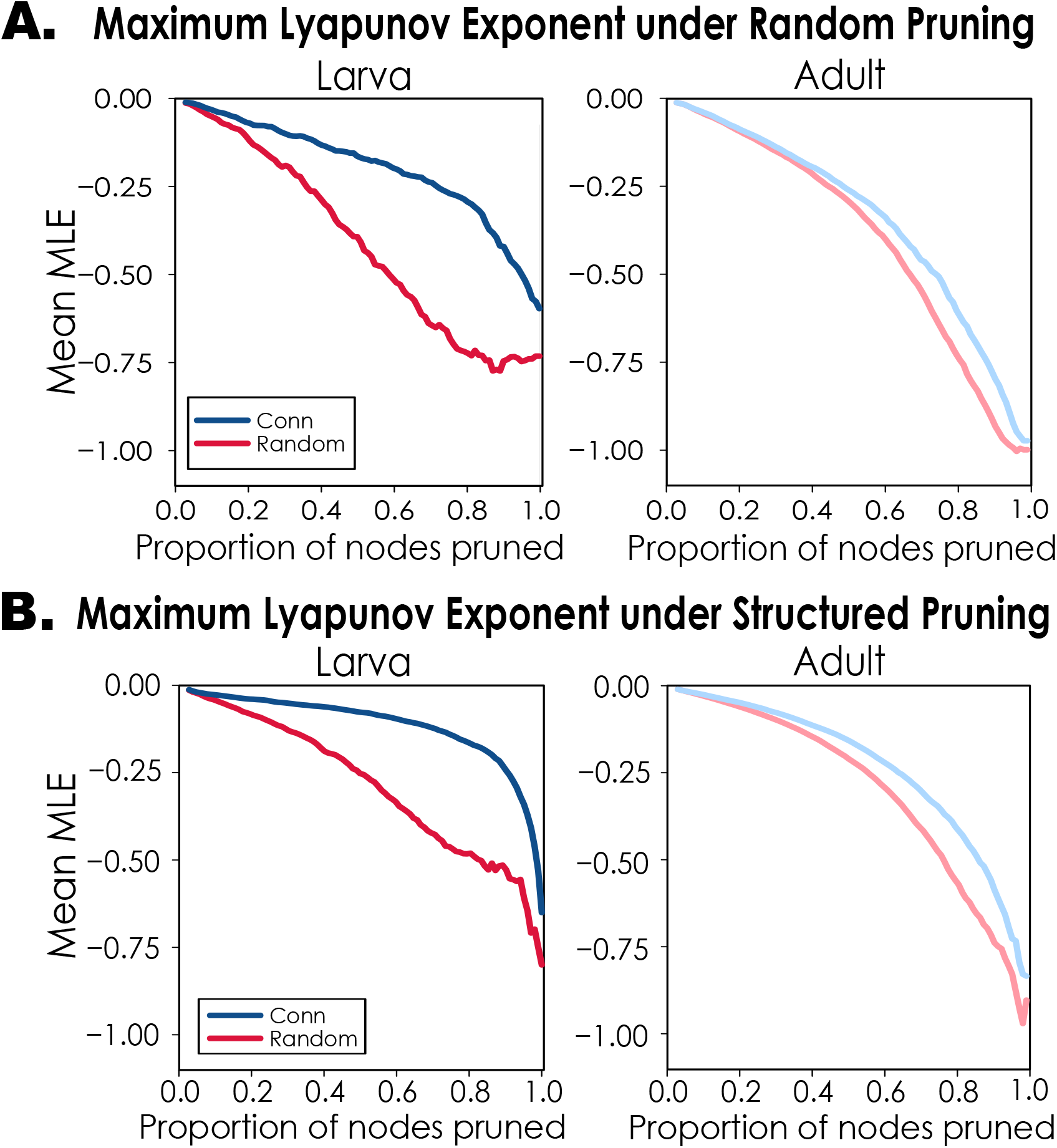
Mean Maximum Lyapunov Exponent of connectome and random ESN dynamics under random (A) and structured pruning.

In the main text we demonstrated how relative spectral radius of connectome-based and random networks decreases under random pruning of neurons. This is closely connected to dynamics, measured here by the mean maximum Lyapunov exponent. Here we show the effect of random and structured (ordered by mean weighted task variance) pruning on mean MLE.

#### Supplementary Note 5: Self-Recurrency and Relative Spectral Radius under Random Pruning

In the main text we analysed networks that contain both sparse off-diagonal connectivity and a controlled fraction of self-connections on the diagonal. Here we derive a simple approximation for the spectral radius that captures the competition between the random bulk spectrum and the diagonal self-recurrency. We first motivate the idea, then derive the approximation for Uniform entries (A), and then provide the equivalent Gaussian case (B).

Consider a random matrix **W** ∈ ℝ*N ×N* with sparsity *S*. A fraction *q* of neurons possess non-zero self-connections.

All non-zero weights are drawn i.i.d. from a zero-mean distribution. The matrix contains two structural components:

- **Diagonal component**: *Nq* self-connections.
- **Off-diagonal component**: the remaining non-zero entries.

The intuition for the derivation can be formalised using the Gershgorin Circle Theorem (9), which states that every eigenvalue *λ* of a matrix **W** lies within at least one Gershgorin disc

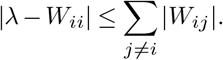

Each disc is therefore centred at the diagonal entry *W*_*ii*_ with radius equal to the absolute row sum of the off-diagonal elements. In sparse random networks without strong diagonal terms, the radii dominate the centres, producing large discs centred near zero and yielding a bulk eigenspectrum determined primarily by the off-diagonal row sums. However, when sufficiently many/large diagonal entries are present, some discs become centred far from the origin. In this regime the spectral radius may be dominated by the diagonal centres rather than the off-diagonal bulk.

**Figure S20.**
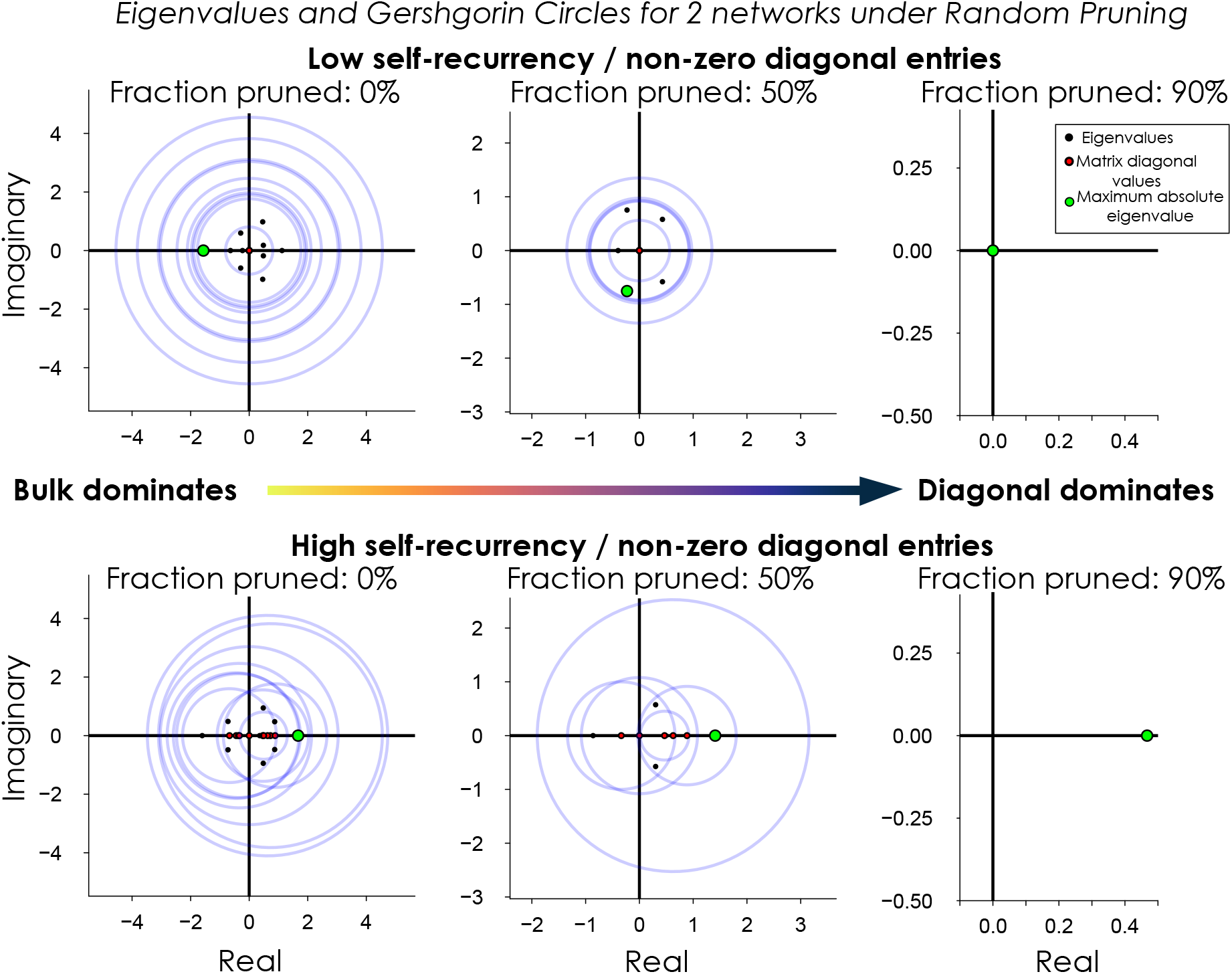
Eigenspectra and Gershgorin radii for two matrices (low self-recurrency – above, high self-recurrency – below) under random pruning (0%, 50%, 90%). As pruning progresses, the eigenspectrum of the high self-recurrent matrix moves from being bulk dominated (left) to being diagonal dominated (right).

Random node pruning reduces the number of off-diagonal connections and therefore shrinks the Gershgorin radii approximately proportionally to (1 − *f*), while the diagonal centres remain unchanged except for the removal of some nodes. Consequently, pruning gradually shifts the dominant contribution to the spectral radius from the bulk row sums toward the surviving diagonal entries.

**Figure** S20 illustrates this transition. Before pruning, the bulk dominates and neither network has an eigenspectrum dominated by the diagonal entries (the red dots on *x* axis). As pruning progresses the radii shrink, and in networks with more diagonal entries and high self-recurrency (bottom row) the circle centres increasingly determine the outermost discs. This allows the self-recurrent network to maintain a larger spectral radius under pruning. Hence we approximate the relative spectral radius by the dominant contribution of the bulk or the diagonal.

##### A. Uniformly Distrib*u*ted Entries

Assume all non-zero weights are drawn i.i.d. from *W*_*ij*_ ∼ 𝒰(−*A, A*).

###### Diagonal contribution

Among the *Nq* diagonal/self-connections, the spectral contribution is governed by the maximum absolute value. Using order statistics for the Uniform distribution (7), the expected maximum absolute value is

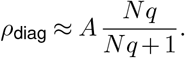

###### Bulk contribution. The off-diagonal entries form a sparse random matrix with *k* non-zero inputs per neuron on average, where *k* (the expected number of off-diagonal non-zeros per row) is

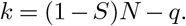

For Uniform weights with variance 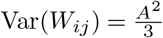 random matrix theory (Supplementary Material 1) predicts that the bulk spectrum scales as

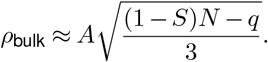

###### Combined approximation

The spectral radius of the full matrix is approximated by the dominant contribution:

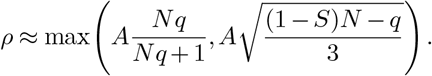

###### Effect of random pruning

When a fraction *f* of neurons is randomly pruned, the size of the new matrix is *N* ^*′*^ = (1 − *f*)*N*. Thus

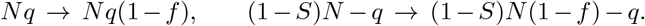

The two spectral contributions become

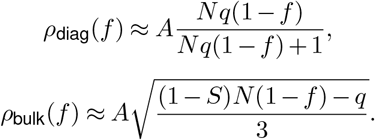

The resulting spectral radius approximation is

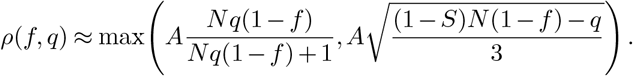

Normalising by the un-pruned, baseline spectral radius

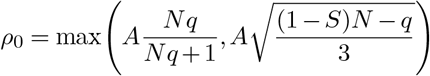

gives the predicted approximate relative spectral radius

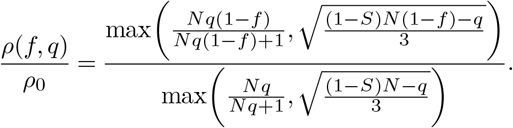

This expression captures the transition between two regimes: when diagonal entries dominate, the spectral radius is governed by extreme value statistics, whereas when the sparse bulk dominates, the relative spectral radius follows the characteristic 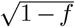 decay of random matrix spectra under node removal.

We note that this theoretical approximation with uniformly distributed weights is limited in that the expected maximum absolute value will saturate at *A* for large *N*, which means that the initial relative diagonal contribution will be capped, and the bulk contribution will dominate, reducing the effectiveness of this theoretical approximation as network size increases. We ran simulations of networks with uniformly distributed weights, sizes *N* = 100, 200 and 300, sparsity *S* = 0.985, varying the fraction of self-recurrent nodes *q* between 0 and 1, and pruning nodes randomly (**Figure** S21). As network size increased, the alignment between the theoretical approximation and simulations diverged. However, both continued to demonstrate that higher self-recurrency yields greater robustness of relative spectral radius to random node deletion. Regardless of the limitation of the theory in larger *N* cases (where the diagonal component saturates), the theoretical approximation still illustrates the important role of increasing self-recurrency in maintaining spectral radius robustness to random pruning, and especially so in relatively small and highly sparse networks.

**Figure S21.**
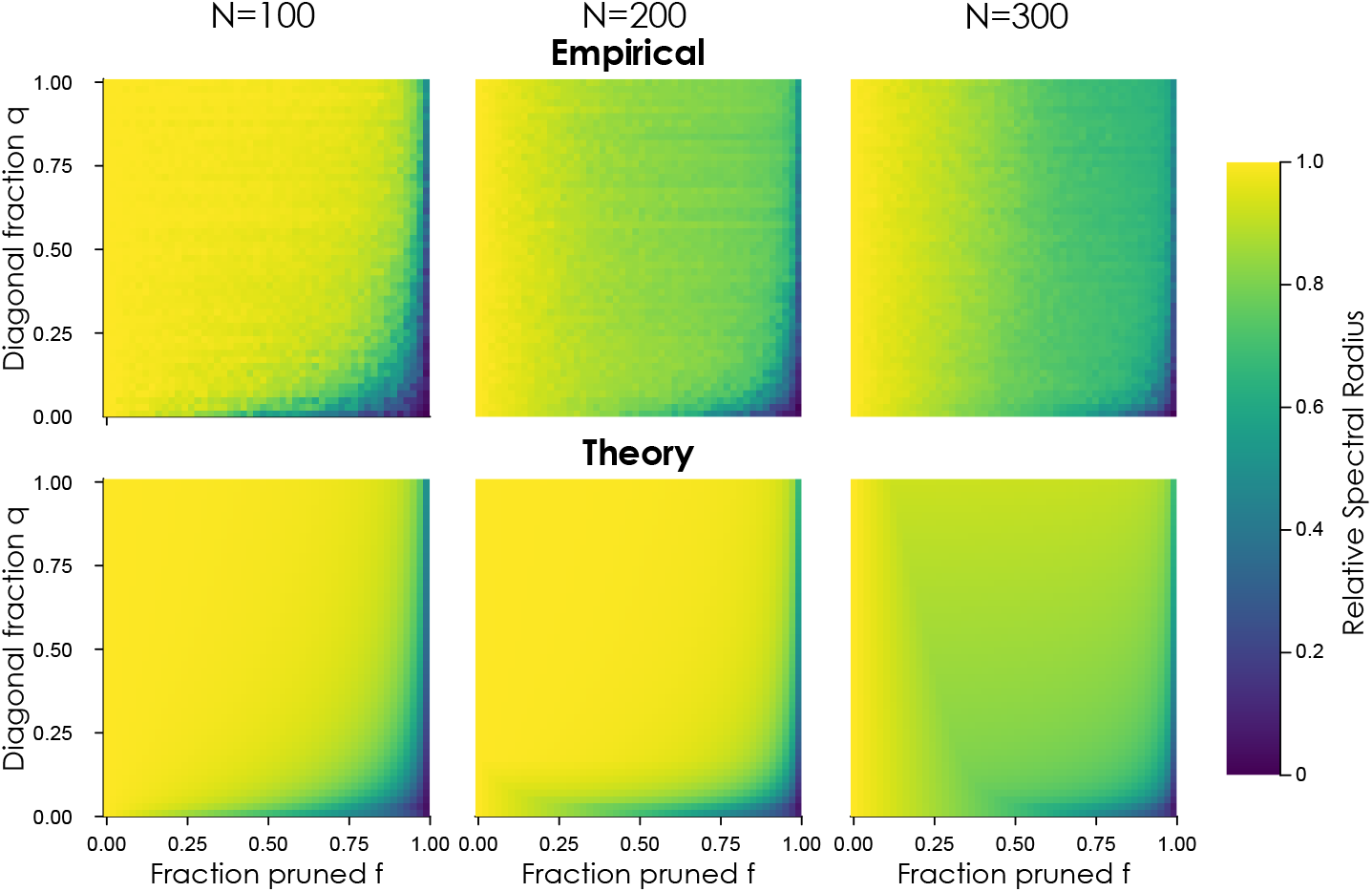
Relative spectral radius under random pruning and varying self-recurrency: Simulation (above) vs theory (below) for uniform weights and different network sizes (*N* = 100, 200, 300 left to right).

One way to address the limitation of the saturating diagonal contribution is to consider weights drawn from a Gaussian distribution. In this case the extreme value statistics grow with network size, allowing the diagonal component to contribute more effectively for larger networks (**Figure** S22). We therefore derive the corresponding Gaussian-weight approximation of the relative spectral radius below.

##### B. Gaussian Distributed Entries

We now assume non-zero weights are drawn from *W*_*ij*_ ∼ 𝒩 (0, *σ*^2^).

###### Diagonal contribution

For *Nq* Gaussian diagonal entries, extreme value theory gives the expected maximum magnitude (15)

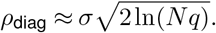

###### Bulk contrib*u*tion

For random matrices with variance *σ*^2^ and *k* non-zero inputs per neuron, the bulk spectrum scales as

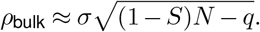

###### Combined approximation

The spectral radius is therefore approximated by

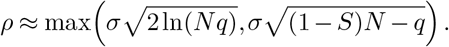

Thus, after pruning a fraction *f* of nodes,

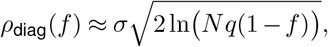

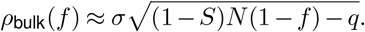

The resulting approximation becomes

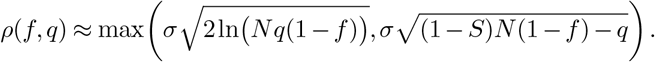

Normalising by the baseline spectral radius

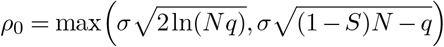

gives the predicted relative spectral radius

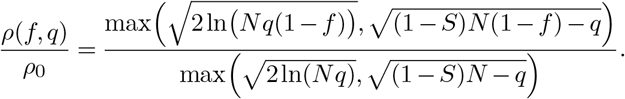

This formulation again highlights the competition between diagonal extreme-value statistics and the bulk random matrix spectrum. Increasing the fraction of self-recurrent neurons increases the likelihood that the diagonal component dominates, thereby stabilising the spectral radius under random pruning.

We ran the same comparison as with the Uniform-weight case for network sizes *N* = 100, 200 and 300, sparsity *S* = 0.985, with Gaussian weights (mean 0 and variance 1), finding that the Gaussian theoretical approximation better tracked the empirical relative spectral radius for increasing network size (**Figure** S22). As in the Uniform-weight case, networks with higher self-recurrency continued to show a greater robustness of spectral radius to random pruning, both in empirical and theoretical results.

**Figure S22.**
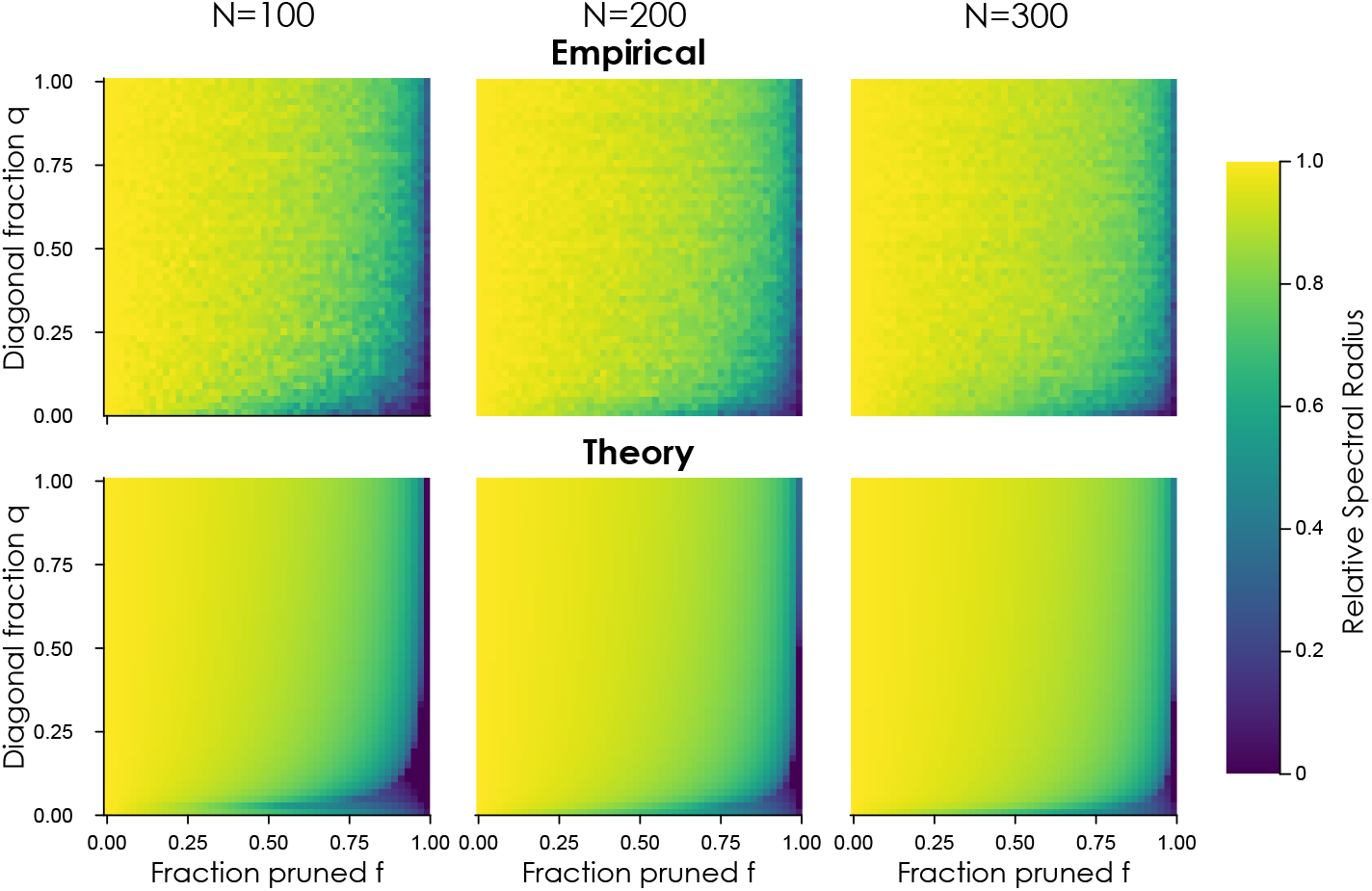
Relative spectral radius under random pr*u*ning and varying self-rec*u*rrency: Simulation (above) vs theory (below) for Gaussian weights and different network sizes (*N* = 100, 200, 300 left to right).

### Supplementary Note 6: The effect of sparsity on dynamics

We observed in the main results that the maximum Lyapunov exponents of the connectome and random networks did not typically exhibit a transition to chaos (MLE > 0) for increasing spectral radius — a transition that is well documented in the literature on random networks (19, 13, 24). To test the relationship between sparsity and dynamics, we calculated the maximum Lyapunov exponent and Lyapunov dimension for random networks (size *N* = 100) with three different sparsity levels (*S* = 0.90, 0.95, 0.985) and found that for lower levels of sparsity (0.9 and 0.95), the transition to chaos is observed in mean intrinsic dynamics (**Figure** S23 left). Furthermore, the networks with lower sparsity also exhibited higher dimensional dynamics (**Figure** S23 right).

**Figure S23.**
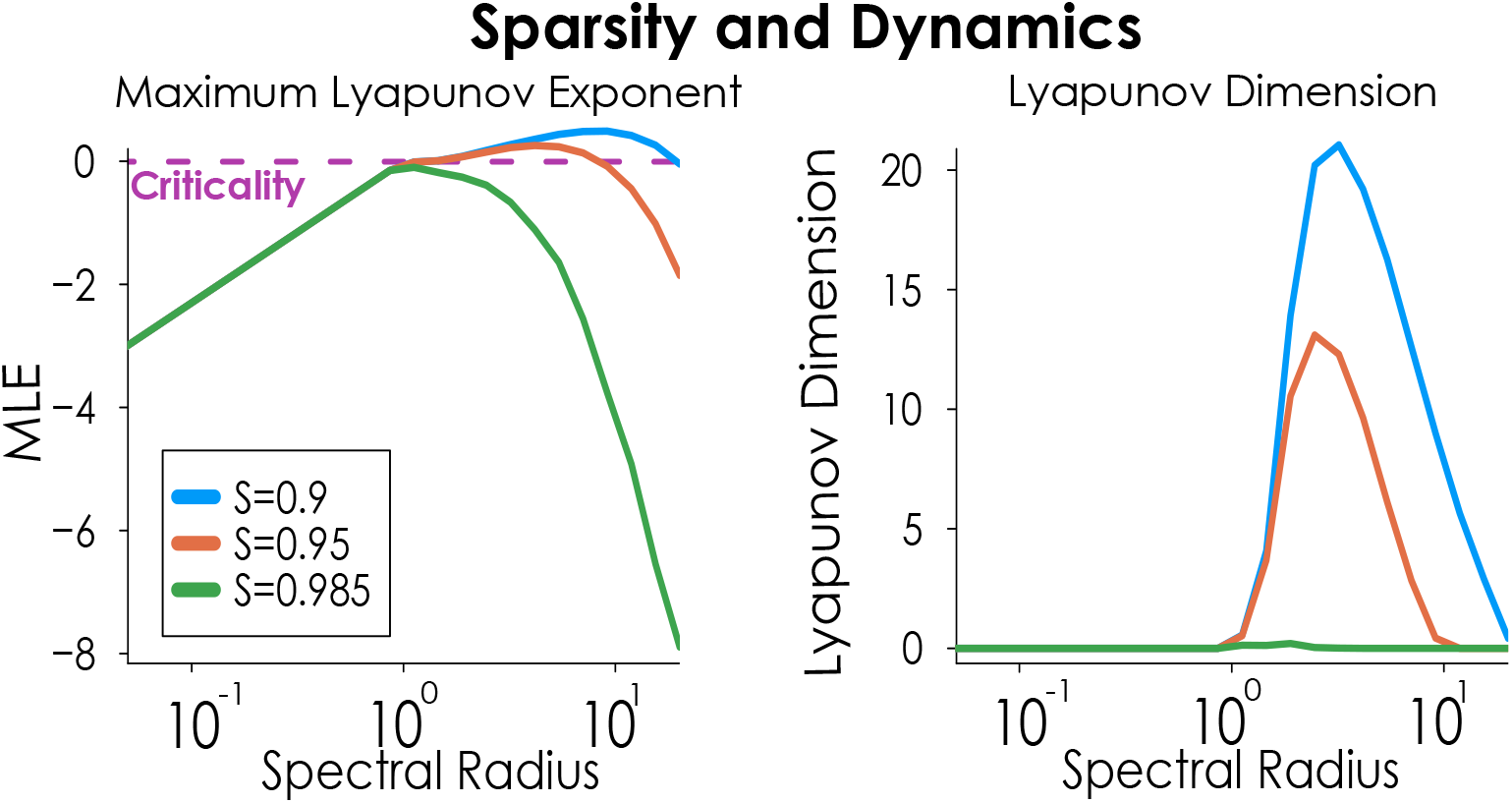
Maximum Lyapunov exponent (left) and Lyapunov dimension (right) across a range of spectral radius values, for different levels of network sparsity (*S* = 0.9, 0.95, 0.985).

### Supplementary Note 7: Network Eigenspectra

**Figure S24.**
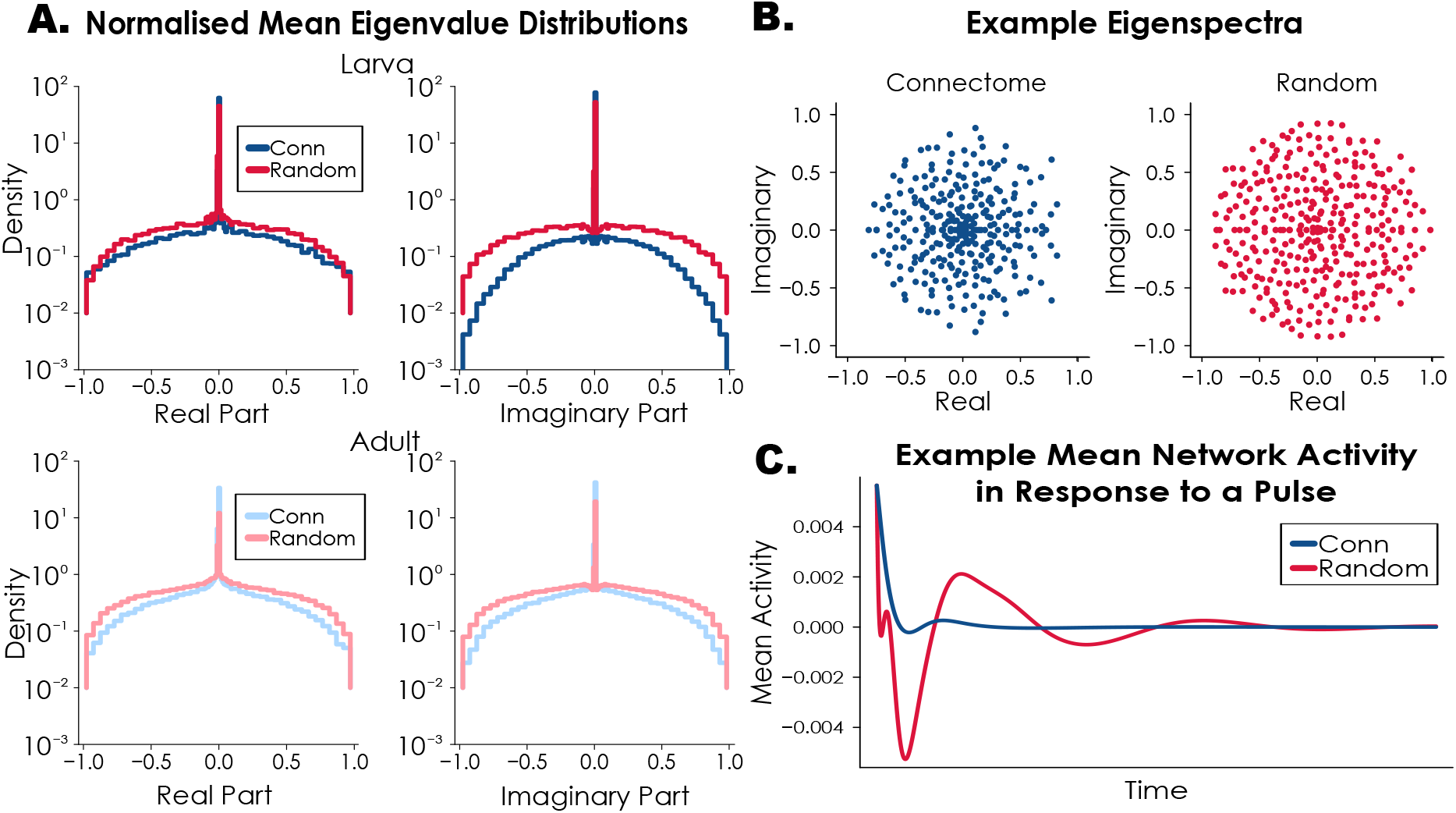
Spectral properties of connectome and random networks. A. The mean distribution of real and imaginary parts of the network eigenvalues. B. Example eigenspectra. C. The example response of networks to a pulse, highlighting the role of eigenvalues on transients and frequencies.

### Supplementary Note 8: Configuration Model Networks

We generated Configuration Model networks as an additional random baseline for comparison with connectome-based networks. These networks are random subject to the constraint that their in- and out-degree distributions match those of the corresponding connectomes. They are structurally closer to the connectome ESNs than Erdoős-Rényi graphs. **Figure** S25 presents select results from the main text with the Configuration Model networks included.

**Figure S25.**
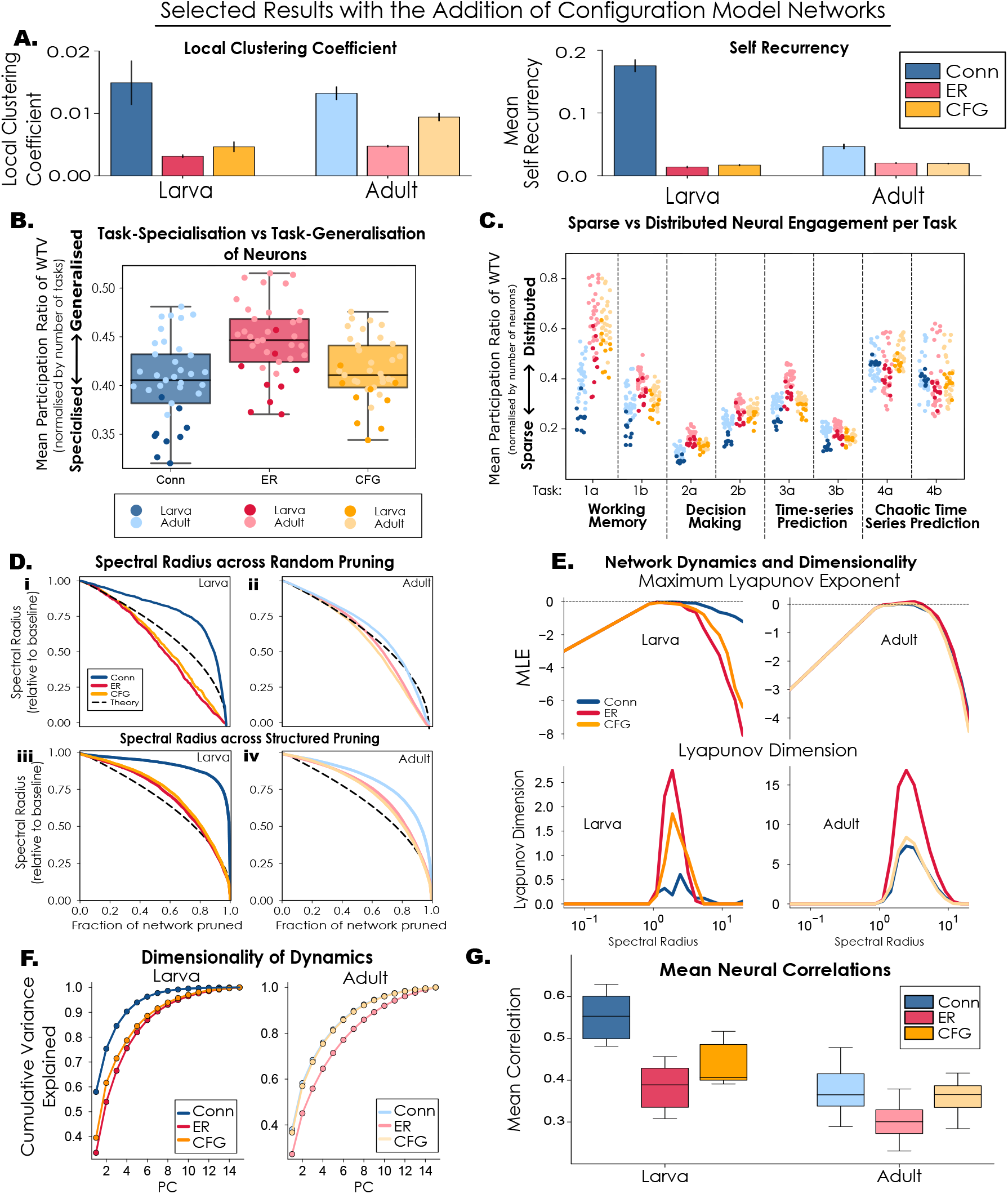
Select res*u*lts from the main paper with Config*u*ration Model networks incl*u*ded. “Conn” = connectome, “ER” = Erdoős-Rényi, “CFG” = Config*u*ration Model. **A**. Mean Local clustering coefficient (left) and Self-recurrency (right). Bars denote standard error of the mean. **B**. Mean participation ratio of weighted task variance for each neuron averaged across 8 tasks. **C**. Mean participation ratio of weighted task variance averaged across networks for each task.**D**. Relative Spectral Radius across Random (above) and Structured (below) neuron pruning. **E**. Network dynamics and dimensionality: Mean Maximum Lyapunov Exponent (above) and Lyapunov Dimension (below).**F**. Dimensionality of Dynamics measured by Cumulative Variance Explained. **G**. Dimensionality of Dynamics measured by Mean Absolute Pairwise Neural Correlations.

### Supplementary Note 9: Cell Types and Weighted Task Variance

**Figure S26.**
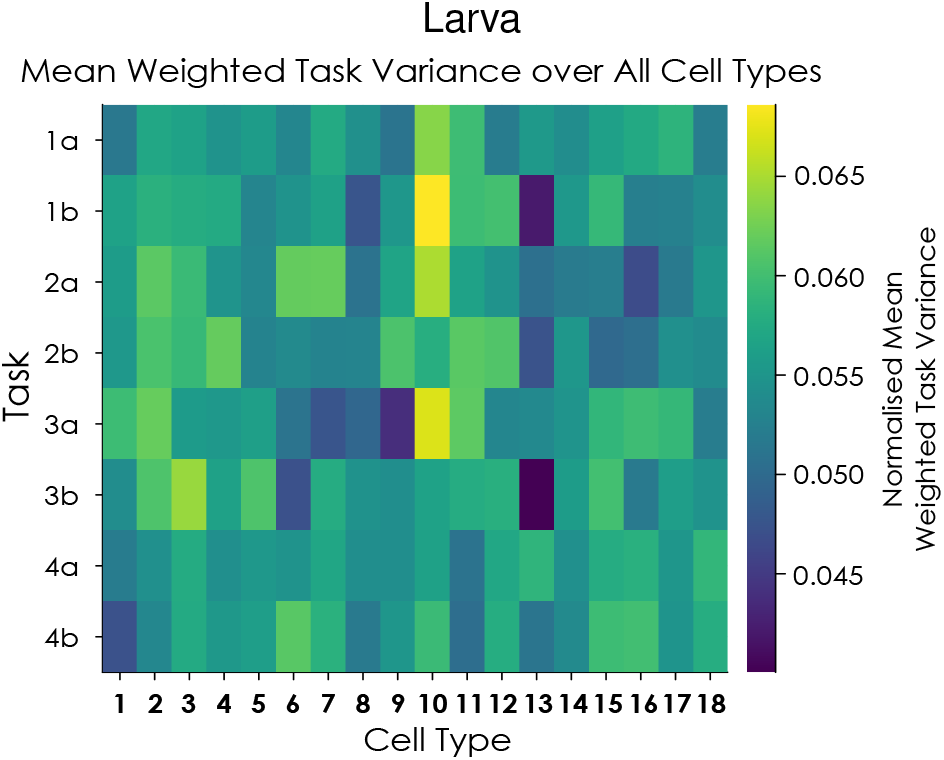
Overall normalised mean weighted task variance for Larva cell types over the 8 Tasks.

**Figure S27.**
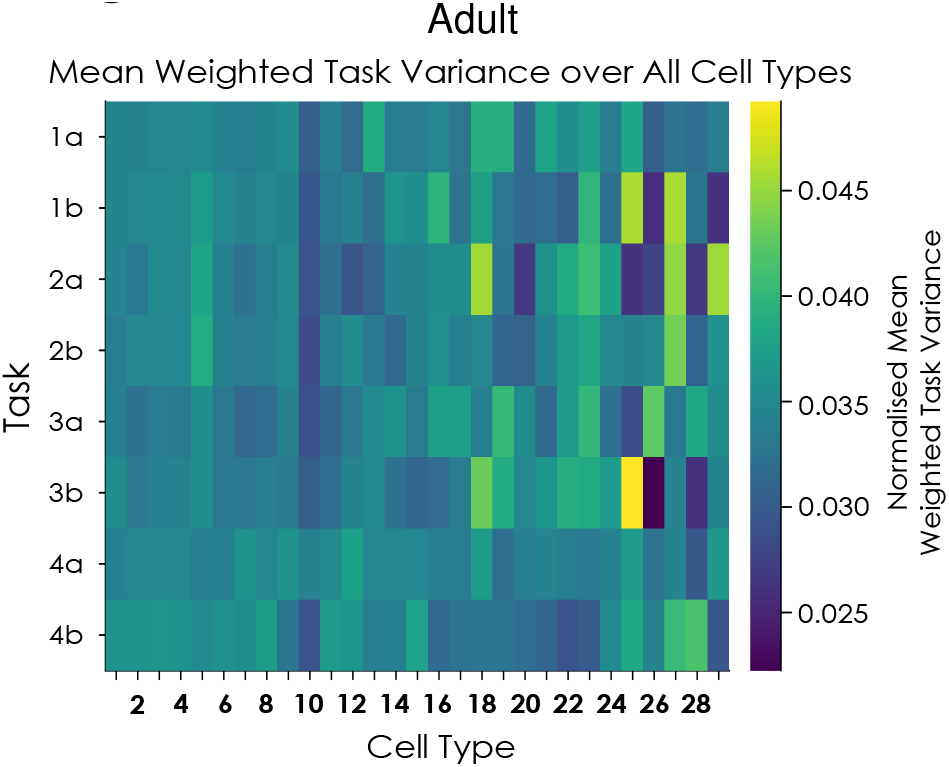
Overall normalised mean weighted task variance for Adult cell types over the 8 Tasks.

#### Larva Cell Types

**1**. Convergence Neurons, **2**. Descending Neurons to Subesophageal Zone, **3**. Descending Neurons to Ventral Nerve Cord, **4**. Kenyon Cells, **5**. Lateral Horn Neurons, **6**. Local Neurons, **7**. Mushroom Body Feedback Neurons, **8**. Mushroom Body Feedforward Neurons, **9**. Mushroom Body Input Neurons, **10**. Mushroom Body Output Neurons, **11**. Projection Neurons, **12**. Somatosensory Projection Neurons, **13**. Axons terminating on Ring Gland Neurons, **14**. Unknown, **15**. Ascending Neurons, **16**. Pre-descending Neurons to Subesophageal Zone, **17**. Pre-descending Neurons to Ventral Nerve Cord, **18**. Sensory Neurons.

#### Adult Cell Types

**1**. Central Complex, **2**. Superior Medial Protocerebrum, **3**. Superior Lateral Protocerebrum, **4**. Lateral Horn, **5**. Superior Intermediate Protocerebrum, **6**. Mushroom Body, **7**. Clamp, **8**. Unknown, **9**. Fruitless, **10**. Descending, **11**. Crepine, **12**. Posterior Ventrolateral Protocerebrum, **13**. Sensory, **14**. Antler, **15**. Anterior Ventrolateral Protocerebrum, **16**. Anterior Optic Tubercle, **17**. Inferior Bridge, **18**. Visual Projection and Optic Lobe, **19**. Posterior Slope, **20**. Lateral Accessory Lobe, **21**. Posterior Lateral Protocerebrum, **22**. Circadian, **23**. Peptidergic, **24**. Antennal Lobe, Vest, **26**. Serotonergic, **27**. Dopaminergic, **28**. Wedge, **29**. Octopaminergic.

